# Instant Prior-Free Resolution Enhancement for Cross-Modality Microscopy

**DOI:** 10.64898/2026.05.28.728601

**Authors:** Haohong Gan, Shiyi Peng, Hailian Hu, Xuan You, Yabo Guo, Ruiyang Guo, Ziliang Chen, Jun Qian

## Abstract

The resolving power of optical microscopy is fundamentally constrained by the diffraction of light, limiting our ability to visualize subcellular structures. Computational methods, particularly deconvolution, can restore blurred images but critically depend on an accurate point spread function (PSF), whose estimation is often impractical and error-prone, leading to artifacts. Here, we introduce Nonlinear Fourier Re-weighting (NFR), a rapid algorithm that operates without any prior knowledge of the imaging system, achieving deconvolution-like effects through a single logarithmic mapping of the image’s Fourier spectrum. This non-iterative process re-balances spatial frequency components to computationally reverse the effects of optical blurring. We demonstrate that NFR robustly enhances resolution beyond the Sparrow limit and recovers authentic structural details. NFR excels where traditional methods fail, remaining effective in the presence of severe optical aberrations and high noise. Furthermore, NFR synergistically improves the output of super-resolution modalities like structured illumination microscopy (SIM), and its near-instantaneous processing enables real-time enhancement of dynamic biological processes, such as in vivo multi-photon microscopic imaging deep within scattering tissue. By decoupling high-fidelity image restoration from system modeling, NFR offers a powerful, accessible, and universally applicable tool for improving image quality across diverse microscopic techniques, facilitating the analysis of large datasets and the discovery of previously obscured biological phenomena.

## INTRODUCTION

The ability to resolve subcellular structures is fundamental to understanding biology^1,2^, making the pursuit of higher resolution a central and enduring theme in optical microscopy. For over a century, the diffraction of light imposed a theoretical limit on the finest detail that could be observed. In recent decades, two principal strategies have emerged to overcome this barrier. The first, leveraging sophisticated hardware and photophysics, gave rise to super-resolution microscopic techniques. These include methods like Structured Illumination Microscopy (SIM)^3^, which modulates the excitation light to achieve frequency shifting, as well as Stimulated Emission Depletion Microscopy (STED)^4^ and Single-Molecule Localization Microscopy (SMLM)^5,6^, which break the diffraction limit by fundamentally altering the fluorescence emission process to achieve nanometer-scale resolution. While transformative, these methods often require specialized instrumentation or complex sample preparation, and can be limited in their application to live-cell imaging^7^ due to phototoxicity and speed constraints.

The second strategy is computational, seeking to maximize the information content captured by the microscope hardware. Within this domain, one family of methods—including Super-resolution Optical Fluctuation Imaging (SOFI)^8,9^ and Super-Resolution Radial Fluctuations (SRRF)^10^—analyzes the statistical fluctuations of fluorophores over a temporal sequence. By trading temporal resolution for spatial information, these techniques can computationally reconstruct an image that surpasses the system’s cutoff frequency, requiring no specialized hardware. Parallel to these fluctuation-based approaches, the most established and broadly applicable software-based method is deconvolution. It aims to reassign photons and reverse the blurring effect of the microscope’s PSF, thereby enhancing both contrast and resolution within the system’s theoretical frequency passband.

However, the efficacy of conventional deconvolution (such as Wiener filtering) ^11,12^ is critically dependent on an accurate model of the PSF. Obtaining this PSF is a major practical bottleneck: it requires meticulous measurement using sub-resolution beads and is sensitive to experimental variations like sample-induced aberrations, imaging depth, and system stability, often leading to suboptimal results or artifacts^13^. To circumvent the need for a measured PSF, blind deconvolution methods^14,15^ were developed. These algorithms attempt to estimate the PSF and the true object simultaneously from the captured image. While powerful, they are invariably iterative, computationally intensive, and consequently slow^16^. This low temporal resolution makes them unsuitable for real-time feedback, live-cell imaging, or high-throughput screening applications. In addition, the iterative estimation process may not converge, and the PSF may be incorrectly estimated, resulting in artifacts^17^. This field is thus faced with a critical question: is it possible to achieve the resolution-enhancing benefits of deconvolution without the need to measure or estimate a PSF and without the cost of iterative computation?

Answering this question requires a deeper understanding of the image restoration problem from the frequency-domain perspective. The resolution of an imaging system is decided by the shape of its optical transfer function (OTF), the Fourier transform of the PSF, not just on the cut-off frequency. Different modalities exhibit characteristic OTF shapes; For instance, the resolution enhancement in the laser scanning confocal microscope stems from its confocal setup and a pinhole that rejects out-of-focus fluorescence^18^. Consequently, the effective PSF of an ideal confocal system can be described as the product of its excitation PSF and detection PSF:

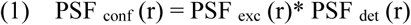

Where **r** represents the spatial coordinate vector. This results in a lateral resolution enhancement of approximately √2 (≈1.4) times that of a conventional wide-field microscope^19^. In the frequency domain, which is described by the spatial frequency vector **k** (the Fourier conjugate of **r**), the OTF of confocal microscope is the Fourier transform of this effective PSF^20^, which translates to the convolution of the excitation and detection OTFs:

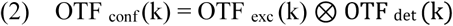

By neglecting the Stokes shift, the two OTFs are identical. As the OTF of wide-field microscope can be approximated as a triangular function^20^, their convolution results in the OTF of confocal microscope with a support that extends to twice the cutoff frequency of wide-field microscope (k _cutoff,conf_ = 2⋅k _cutoff,WF_). Interestingly, SIM also achieves a doubled cutoff frequency (k _cutoff,SIM_ = 2⋅k _cutoff,WF_) by synthetically extending the OTF’s support^3^, yet it yields a genuine two-fold resolution improvement. We are thus faced with two distinct modalities, confocal and SIM, that share the same theoretical OTF cutoff frequency yet deliver different effective resolutions (∼1.4× vs. 2×). This discrepancy arises because the high-frequency components of the OTF of confocal microscope, shaped by OTF convolution, are far more attenuated than those of the synthetically assembled OTF of SIM. This comparison powerfully corroborates the principle that the OTF’s shape, not merely its cutoff frequency, is the true determinant of a microscope’s resolving power. This also means that confocal microscope and SIM have almost the same resolution limit^15,21^. This shared limit is also the conceptual foundation for technologies like Image Scanning Microscopy (ISM)^22^. ISM uses an array detector to dramatically improve the signal-to-noise ratio and then applies deconvolution to reshape the effective OTF, thereby unlocking the confocal system’s latent potential to surpass the two-fold resolution enhancement.

If disparate high-frequency attenuation between confocal microscope and SIM is the key, could resolution be recovered by simply amplifying these frequencies after acquisition? The answer is negative. Critically, a high-quality, “healthy” biological image inherently contains a balanced distribution of energy across both low and high spatial frequencies. Simple post-processing like sharpening, which arbitrarily amplifies high frequencies, fails to restore true resolution and often introduces noise and artifacts. The above perspective clarifies the fundamental nature of deconvolution: it is essentially a re-equalization of the image’s frequency energy.

An intriguing parallel can be drawn from the field of digital imaging. Image sensors typically have a linear response to photon flux, meaning that bright signals occupy a proportionately large portion of the available dynamic range. The human visual system, in contrast, perceives brightness non-linearly, in a manner closer to a logarithmic scale^23,24^; a change from intensity level 0 to 1 can be as perceptible as a change from 127 to 255. To reconcile this discrepancy and optimize the use of bit-depth, techniques like gamma correction^25^ or logarithmic mapping^26^ are routinely applied. These methods remap intensity values to create a more perceptually uniform distribution, effectively re-balancing the dynamic range across different brightness levels. This re-balancing of the intensity domain via a logarithmic transform shares a striking conceptual similarity with the re-balancing of the frequency domain by deconvolution. Both aim to correct a systemic imbalance—one in perceived brightness, the other in spectral power. This analogy provokes a pivotal question: could the principle of logarithmic correction be transposed from the intensity domain to the frequency domain? If so, we might directly re-balance the image’s Fourier spectrum to counteract OTF attenuation, thereby achieving a deconvolution-like enhancement of resolution, but entirely bypassing the need for PSF estimation and iterative processing.

In this work, we introduce Nonlinear Fourier Reweighting (NFR), a novel algorithm that realizes this very principle. NFR directly transposes the concept of logarithmic correction from the intensity domain to the frequency domain, aiming to re-balance the spectral power between attenuated high frequencies and dominant low frequencies. Without relying on any system priors or iterative optimization, NFR achieves a significant resolution enhancement, recovering structural details that surpass the Sparrow criterion for diffraction-limited systems. The entire process is accomplished in a single computational step, requiring only one Fast Fourier Transform (FFT) and its inverse. We demonstrate the robustness and versatility of NFR across a wide range of imaging modalities—including wide-field, confocal, and multi-photon microscope—and for various data dimensionalities, such as 1D signals, 2D images, 3D stacks, and time-series. Crucially, it performs effectively even on images with low signal-to-noise ratios. Its freedom from PSF dependencies, combined with near-instantaneous processing speeds, establishes NFR as a powerful and practical tool for real-time image quality enhancement, high-throughput analysis of large datasets, and the discovery of subtle biological structures previously obscured by system blur.

## RESULTS

### Principle of NFR: A Prior-free approach for spectral re-balancing

Conventional deconvolution methods, such as the Richardson-Lucy (RL) algorithm^27,28^, can significantly enhance image resolution. However, they suffer from two major limitations: they depend critically on an accurate model of the PSF, and their iterative nature makes them computationally expensive and prone to amplifying noise and generating artifacts, especially when the PSF is mismatched.

To circumvent these limitations, we re-frame the goal of image restoration not as an inverse problem, but as a task of spectral re-balancing—restoring the energy balance between high and low spatial frequencies. We draw inspiration from a conceptually related problem in camera Image Signal Processor (ISP) design, where logarithmic mapping is used to compress a wide dynamic range of light intensities into a limited bit depth. We translated this principle to the frequency domain. Our NFR algorithm applies a non-linear logarithmic re-weighting (Fig. 1a), directly to the Fourier amplitude of the image:

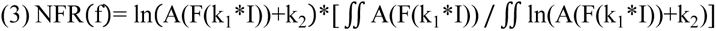

**Fig. 1.**
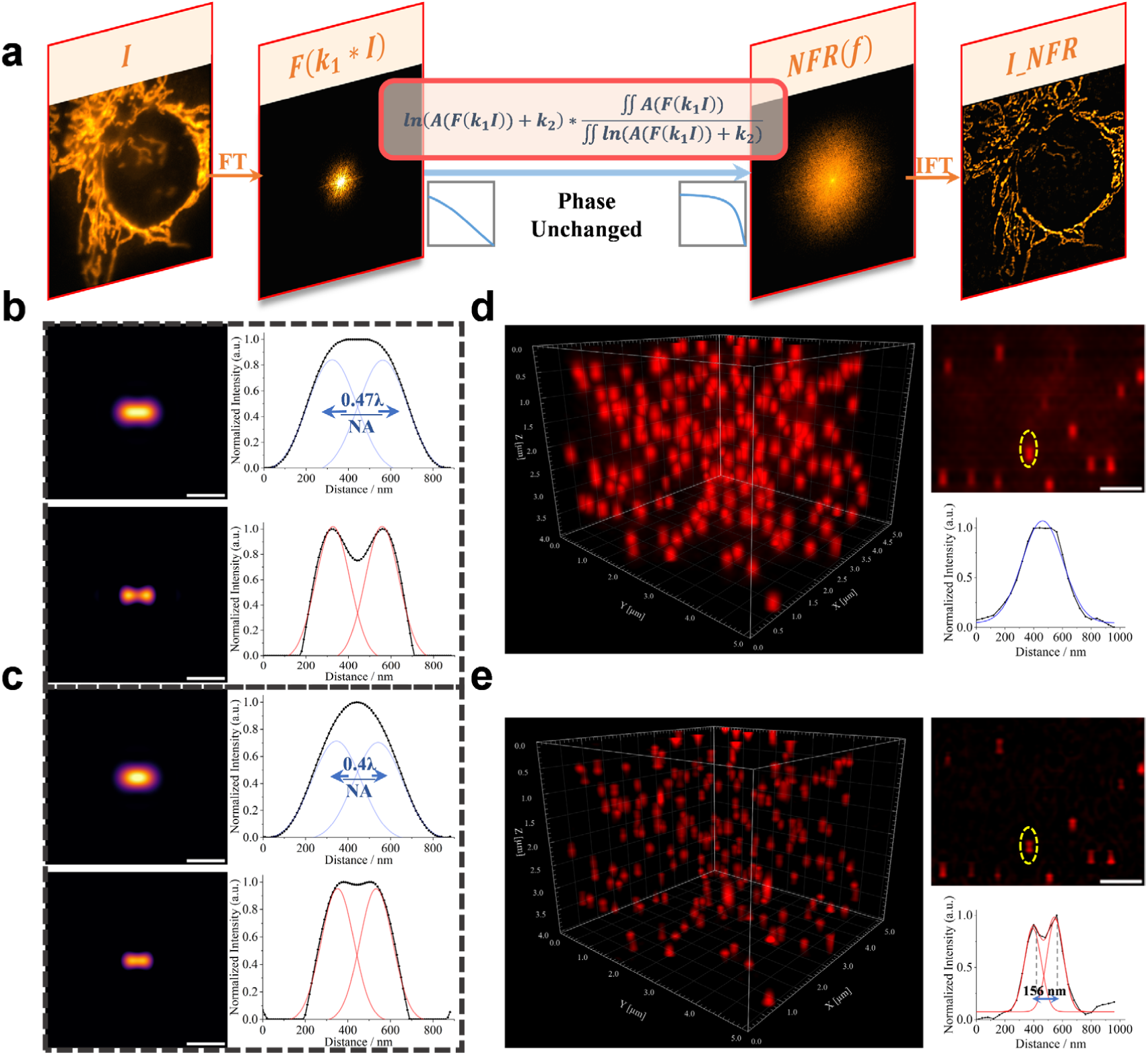
Principle and simulated performance of the NFR algorithm. **a**, Schematic of the NFR workflow for image enhancement. **b, c** Demonstration of lateral resolution enhancement beyond the Sparrow criterion (≈0.47λ/NA). **b,** Two point sources separated by 0.47λ/NA. The diffraction-limited image (top) shows them as unresolved, while the NFR-processed image (bottom) clearly separates them. **c**, The separation is reduced to 0.4λ/NA, well below the Sparrow limit. NFR still resolves the two points. Simulation parameters: λ = 500 nm, NA = 1.0, pixel size = 10 nm. Scale bar, 500 nm. **d, e** Demonstration of axial resolution enhancement. Simulated axial view of two point sources separated by 156 nm before (**d**) and after (**e**) NFR processing. Simulation parameters: λ = 520 nm, NA = 1.4, voxel size = 40 × 40 × 40 nm. Scale bar, 1μm.

This operation effectively transforms the typical, quasi-linear decay of the OTF into a more uniform spectral profile within the system’s passband. This re-balancing directly enhances resolution in a single computational step, requiring only one forward and one inverse Fourier transform with appropriate parameters (Supplementary Figure 1 and Methods).

We validated this principle through simulations. NFR successfully resolves two point-sources separated by a distance smaller than the Sparrow criterion for incoherent imaging, a task where the diffraction-limited system fails. Crucially, as it involves no frequency-domain division, this is achieved without introducing the ringing artifacts common to other sharpening filters (Fig. 1b, c). Importantly, when applied to images of randomly distributed point sources (Supplementary Figure 2), NFR enhances existing structures without fabricating spurious high-frequency details. The method’s effectiveness is not limited to lateral enhancement: by applying the process to 3D stacks, NFR also achieves significant axial resolution enhancement. As demonstrated in simulations, NFR significantly improves axial resolution, enabling the clear separation of two points spaced just 156 nm apart axially—a task impossible for the original diffraction-limited system (Fig. 1d, e).

### Prior-free NFR enhances 2D resolution with high structural fidelity

A central challenge in traditional deconvolution stems from its nature as an inverse problem involving frequency-domain division. Even minute inaccuracies in the PSF model, which are unavoidable in complex biological environments, can lead to division by near-zero values, generating severe artifacts. While advanced algorithms leveraging priors like sparsity (Sparse Deconvolution^14^) or multi-resolution analysis (MRA^29^) have been developed to enhance fluorescence microscopy, they do not escape this fundamental vulnerability. Despite offering potentially higher resolution ceilings or improved noise resilience, they are, without exception, refinements of iterative blind deconvolution frameworks. Consequently, they are still encumbered by initial parameterization and time-consuming iterative computations. In stark contrast, NFR introduces a fundamentally different, prior-free and non-iterative paradigm.

To benchmark NFR against these state-of-the-art methods, we performed direct comparisons on wide-field microscopic images of canonical yet challenging biological structures—microtubules and mitochondrial inner membranes in HeLa cells—using super-resolution SIM images as the ground truth. In densely populated regions of the microtubule network, NFR demonstrated superior resolving power and exhibited the highest structural similarity (SSIM) to the SIM ground truth (Fig. 2a, b). This high-fidelity performance was mirrored in the mitochondrial imaging. While both MRA and NFR successfully resolved two closely-spaced inner cristae that were indistinguishable by other methods, NFR achieved a notably higher degree of separation and again attained the highest SSIM (Fig. 2c, d).

**Fig. 2.**
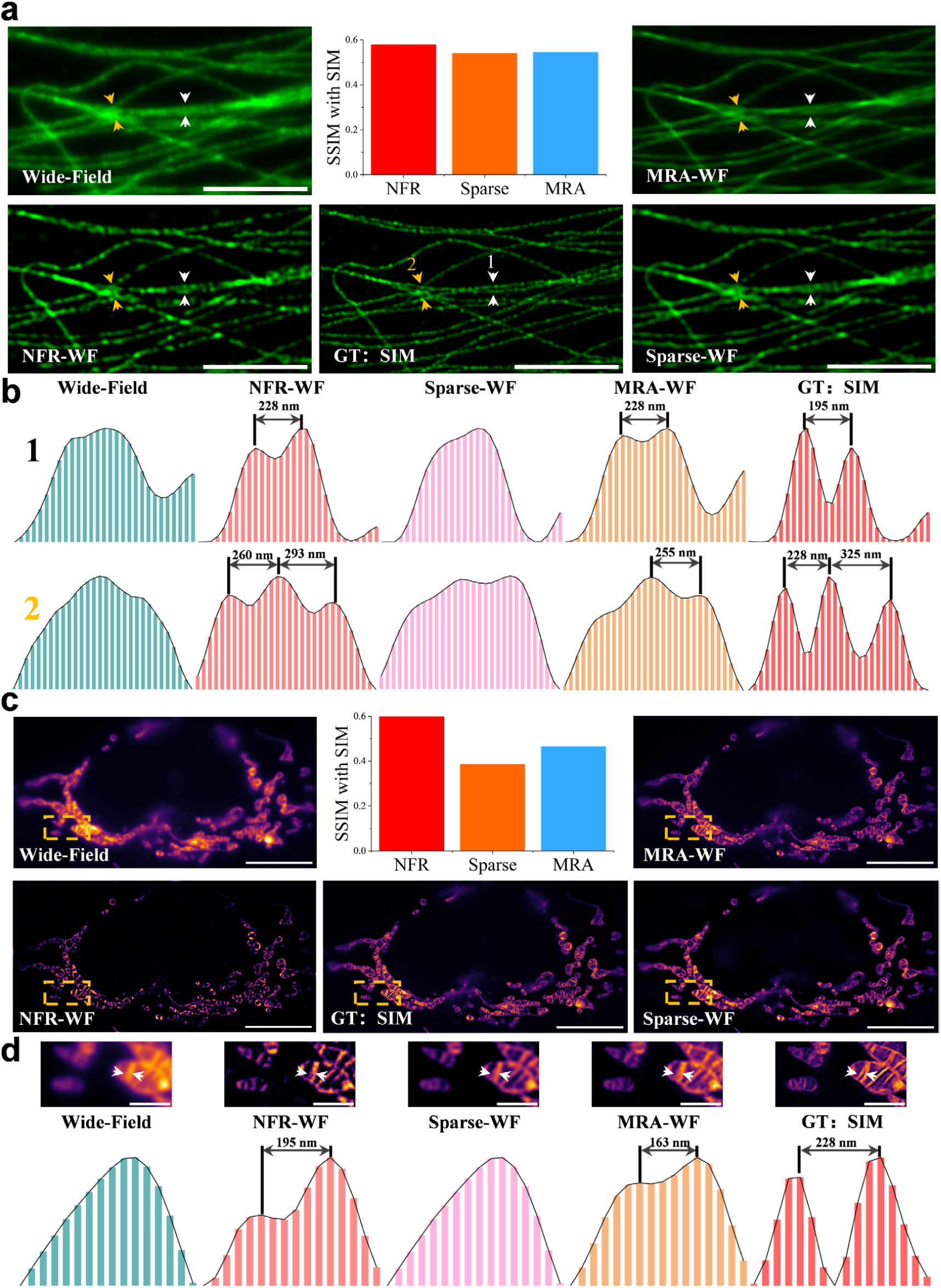
Model-independent NFR enhances resolution while preserving structural fidelity. **a**, Comparison of a wide-field microscopic image of microtubules with its super-resolution SIM (ground truth) counterpart, and the results after enhancement with NFR, MRA, and Sparse Deconvolution. The Structural Similarity Index (SSIM) was calculated against the SIM ground truth, with NFR achieving the highest score (0.58). Scale bar, 5 μm. **b**, Quantitative analysis of the regions indicated in **a**. Normalized intensity profiles across two closely spaced microtubules (region 1) show that both NFR and MRA can resolve the 195 nm gap identified by SIM. In the more complex intersection (region 2), which SIM resolves as three distinct peaks, only NFR successfully separates all three peaks, matching the ground truth with only a single-pixel deviation. **c**, Comparison of a wide-field microscopic image of the mitochondrial inner membrane with its SIM ground truth and corresponding enhanced results. NFR again yields the highest SSIM (0.62). Scale bar, 10 μm. **d**, Magnified views of the mitochondrion indicated by the orange box in **c**. SIM resolves two inner cristae separated by 228 nm. While Sparse Deconvolution fails to resolve them, both MRA and NFR succeed. Notably, NFR exhibits superior separation and fidelity compared to MRA (resolved separation of 195 nm vs. 163 nm, respectively). Scale bar, 2 μm.

Furthermore, while advanced algorithms like Sparse Deconvolution and MRA may achieve competitive results in specific scenarios, their performance is critically dependent on meticulous, trial-and-error parameter tuning (e.g., sparsity, wavelet thresholds). Coupled with their iterative nature, this makes optimization exceedingly time-consuming. NFR’s non-iterative design completely bypasses this bottleneck, enabling real-time parameter adjustments with instantaneous visual feedback and thus offering an unparalleled advantage in speed and usability (Supplementary Video 1). This user-friendly parameter tuning directly translates to robust, high-throughput processing, as the optimized parameters can be universally applied to enhance entire time-lapse sequences, revealing fine subcellular dynamics previously obscured by diffraction (Supplementary Video 2).

Finally, we demonstrated NFR’s versatility, showing it can substantially narrow the performance gap between conventional confocal and Airyscan imaging using the same high-NA objective (Extended Data Fig. 2a), while also enhancing the resolution of super-resolution SIM images (Extended Data Fig. 2b). Our quantitative analyses therefore confirm that NFR is a potent post-processing tool that delivers high-fidelity resolution enhancement without the need for system modeling or iterative computation.

### NFR enhances volumetric resolution in 3D microscopy

Extending image restoration to three dimensions presents substantial challenges for conventional deconvolution. The computational complexity increases by orders of magnitude for 3D stacks. Furthermore, the assumption of a spatially invariant PSF becomes increasingly fragile, particularly in scattering media where the PSF can vary significantly with depth^30^. Accurately measuring or estimating the 3D PSF, especially its axial component, is often intractable. NFR elegantly circumvents these issues. By simply replacing the 2D FFT with its 3D counterpart, the NFR workflow directly extends from two to three dimensions (Fig. 3a), enabling volumetric resolution enhancement without the need for a 3D PSF model.

**Fig. 3.**
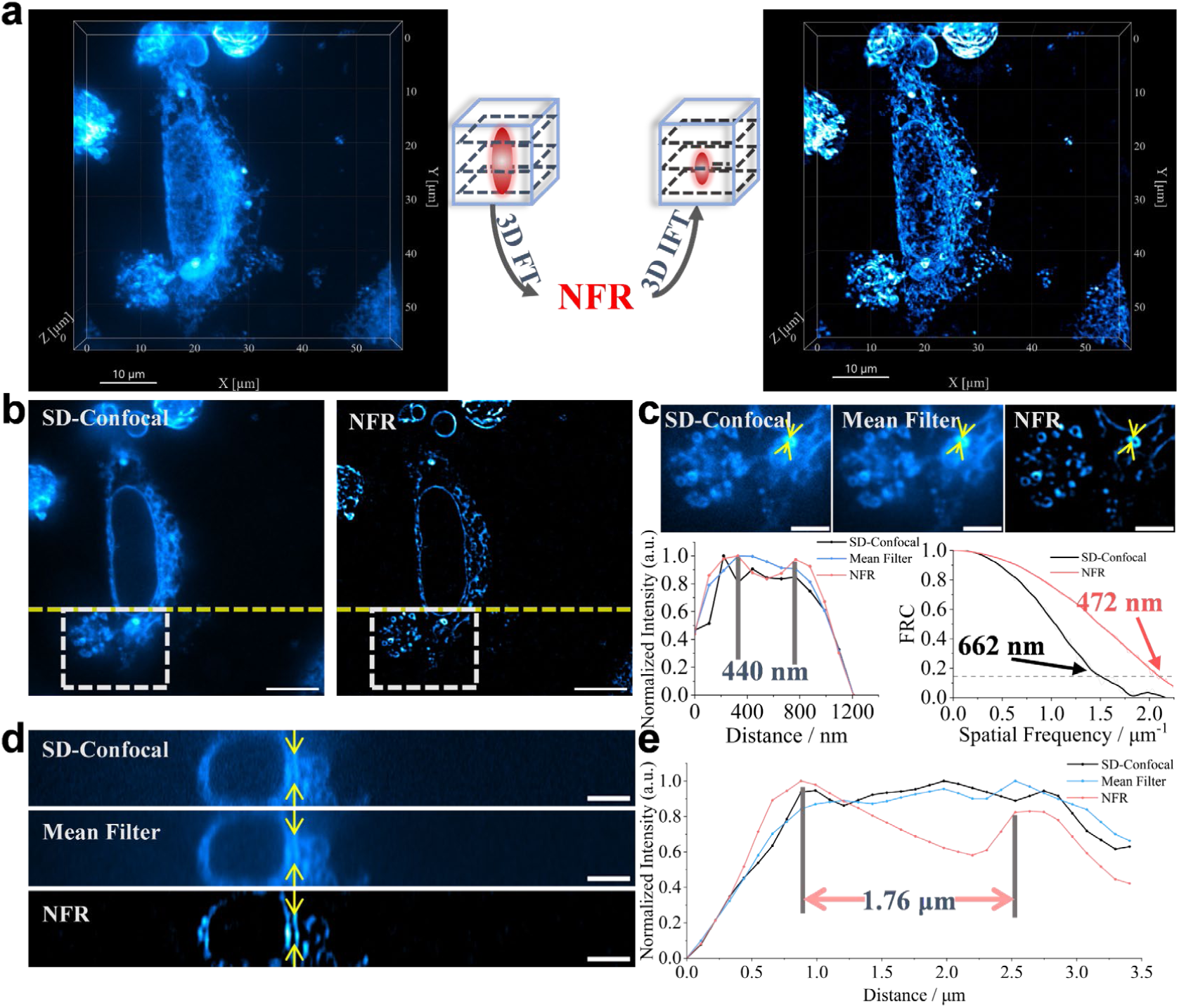
NFR enhances 3D resolution of spinning-disk confocal (SD-confocal) stacks. **a**, 3D volume rendering of an Endoplasmic Reticulum (ER) network in an eRFP-expressing HeLa cell, imaged by SD-confocal microscopy and processed with the 3D NFR workflow. **b**, A representative XY slice from the 3D stack. Scale bar, 10 μm. **c**, Magnified view of the boxed regions in **b**, comparing the raw SD confocal data, a mean-filtered result, and the NFR-processed image. NFR clearly resolves the ring-like cross-sections of ER tubules (∼440 nm separation, yellow arrows). FRC analysis confirms resolution enhancement from 662 nm to 472 nm. Scale bar, 4 μm. **d**, An axial cross-section along the yellow dashed line in **b**, comparing profiles from the raw, mean-filtered, and NFR-processed data (top to bottom). Scale bar, 4 μm. **e**, Quantitative axial intensity profiles of yellow arrows in **d**, showing NFR resolves two structures separated by 1.76 µm that are unresolved in raw and mean-filtered data.

To demonstrate this capability, we applied the 3D NFR algorithm to a publicly volumetric dataset of the Endoplasmic Reticulum (ER) in a HeLa cell^31^, acquired with a spinning-disk confocal (SD-confocal) microscope. In lateral XY slices, NFR processing reveals fine, ring-like cross-sections of ER tubules that are obscured in the raw data and remain unresolved after simple mean filtering (Fig. 3b). This visual improvement is quantified by FRC analysis, which indicates a ∼1.4-fold enhancement (from 662 nm to 472 nm) in lateral resolution compared to the raw data (Fig. 3c). The benefits of NFR extend to the axial dimension. In an axial cross-section, NFR successfully resolves two structures separated by 1.76 μm that are merged into a single feature in both the raw and mean-filtered data (Fig. 3d).

The robustness of 3D NFR is further highlighted in its application to imaging in highly scattering media. When applied to a 3D two-photon microscopic dataset of chloroplasts within a live plant leaf, NFR also provides a marked enhancement in axial resolution, demonstrating its utility in highly scattering biological samples (Extended Data Fig. 3).

### NFR enables high-speed resolution enhancement in dynamic deep live-tissue imaging

A key advantage of NFR’s single-step nature is its computational efficiency, which translates directly to high temporal resolution for post-processing—a critical factor for dynamic live-cell imaging (Extended Data Fig. 4, Supplementary Video 3 and Supplementary Video 4). We then demonstrate this by imaging chloroplast autofluorescence^32^ in a live Arabidopsis thaliana leaf using two-photon microscopy, a sample characterized by low signal fluctuation. We benchmarked NFR against SOFI, a true super-resolution technique that extends the frequency passband by analyzing temporal correlations from fluorescence blinking. While powerful, SOFI’s performance is limited for fluorophores with low fluctuation rates^33^, such as the autofluorescence from chloroplasts.

Using a resonant scanner for high-speed acquisition, we compared three processing pipelines on a time-series acquired with 950 nm excitation: first-order Fourier-SOFI (fSOFI)^34^ on 100 frames (which is equivalent to a multi-frame average), NFR applied to the first-order fSOFI result, and second-order fSOFI on 300 frames (Fig. 4a). While first-order fSOFI significantly improved the SNR, it offered no resolution gain (Fig. 4b). In stark contrast, applying NFR to this averaged result clearly resolved the fine grana structures within the chloroplasts. This NFR-enhanced result was visibly superior to the second-order fSOFI image, which required three times more frames to compute.

**Fig. 4.**
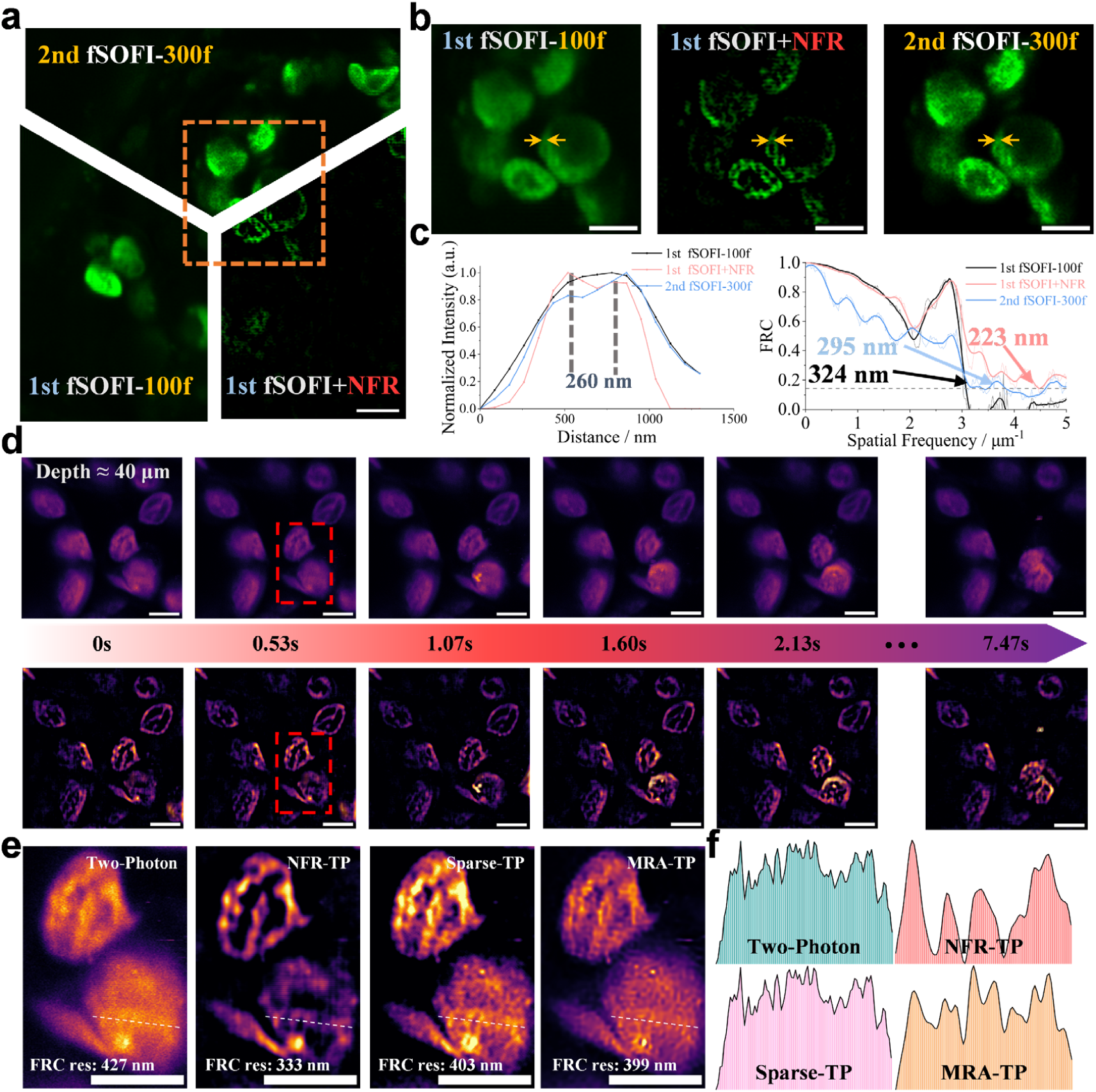
NFR enables high-speed, high-resolution two-photon microscopic imaging in highly scattering live tissue. **a,** Comparison of first-order fSOFI (100 frames), fSOFI combined with NFR (100 frames), and second-order fSOFI (300 frames). The NFR combination reveals fine chloroplast structures not visible with the other methods. Scale bar, 5 μm. **b,** Magnified view of the region indicated by the orange box in **a**. Scale bar, 4 μm. **c,** Intensity profiles of the chloroplast structures indicated by the yellow arrows in **b**, demonstrating that fSOFI+NFR resolves two adjacent structures. Corresponding FRC analysis confirms the superior performance of the NFR-enhanced fSOFI result (223 nm) compared to standard second-order SOFI (295 nm). **d,** High-speed image sequence (3.7 FPS) capturing the photodamage of chloroplasts under high-power excitation. NFR is applied to each frame, demonstrating its compatibility with high-speed acquisition. Scale bar, 5 μm. **e**, Comparison of different resolution enhancement methods applied to the region indicated by the red box in **d**. The raw two-photon microscopic image was processed with NFR, Sparse, and MRA. NFR processing yields the most significant resolution improvement (from 427 nm to 333 nm) while introducing the fewest honeycomb-like artifacts. Scale bar, 5 μm. **f**, Intensity profiles along the white dashed lines in **e**. The NFR profile exhibits the lowest noise and artifact levels.

Quantitative analysis confirms this superiority. Intensity profiles across the grana (indicated by yellow arrows in Fig. 4b) demonstrate that the NFR-enhanced fSOFI result combination successfully resolves two adjacent structures separated by only 260 nm (Fig. 4c). This is corroborated by FRC analysis, which yields an effective resolution of 223 nm for the NFR-enhanced fSOFI result, a significant improvement over both second-order SOFI (295 nm) and the initial first-order fSOFI result (324 nm).

To further demonstrate NFR’s dual advantages in temporal resolution and robustness against scattering, we monitored photodamage in chloroplasts^35^ deep (∼40 μm) within a living Arabidopsis leaf. A time-series was acquired at 3.7 FPS using resonant scanning based two-photon microscope under high-power 850 nm excitation. Applying NFR frame-by-frame provided real-time enhancement of the dynamic process, allowing us to observe the evolving grana structure (Fig. 4d), including the structural changes of internal grana and the gradual fusion of two adjacent chloroplasts (highlighted by red dashed circles in Fig.4d and Supplementary Video 5). We also benchmarked NFR against leading deconvolution algorithms in this challenging, highly scattering environment. Two-photon microscopic imaging in plant leaves inherently suffers from significant scattered background noise. As shown in Fig. 4e, NFR simultaneously enhances resolution and effectively suppresses this background. Crucially, it does so without introducing the prominent honeycomb-like artifacts that often plague iterative deconvolution methods like Sparse and MRA, a conclusion further supported by the intensity profiles (Fig. 4f). This experiment highlights NFR’s capacity to maintain high performance deep within highly scattering tissue, proving that its temporal resolution is ultimately limited only by the microscope’s acquisition speed.

To push the boundaries of NFR’s performance in low-photon, deep-tissue contexts, we applied it to a publicly available three-photon microscopy dataset of YFP-H transgenic mouse medial prefrontal cortex (mPFC)^36^. We focused on a deep volume spanning 600-1600 µm. Given the extremely low photon counts characteristic of three-photon microscopic imaging, we employed a combined NFR and mean filtering strategy to first enhance structural details and then improve the overall SNR. Due to the dataset’s axial undersampling, NFR was applied on a 2D slice-by-slice basis. At depths exceeding 1 mm (∼1050 µm and ∼1475 µm), NFR successfully revealed fine intracellular structures within neurons that were completely absent in both the raw data and the mean-filtered data. This demonstrates that NFR is not merely a filter but a true resolution enhancer, capable of extracting meaningful biological information even under the most challenging signal-starved and high-scattering conditions (Extended Data Fig. 5).

Analogously, the frame-by-frame strategy used for time-series analysis is directly applicable to multi-channel imaging. The NFR algorithm is applied independently to each color channel, and the resulting enhanced channels are then merged to form the final composite image (Supplementary Figure 3). This inherent channel- and frame-independence makes NFR ideally suited for the automated, high-throughput batch processing of large image datasets acquired under consistent conditions.

### NFR as a general method for enhancing resolution

The applicability of the NFR principle extends beyond microscopy. Deconvolution is a cornerstone of astronomical image processing, used to reveal fine structures in distant galaxies or resolve close binary star systems. To demonstrate its broader utility, we applied NFR to an image of the M66 spiral galaxy captured with a 150 mm Newtonian telescope. The algorithm effectively enhanced the contrast and detail of the galaxy’s spiral arms, showcasing its potential for astronomical applications (Fig. 5a).

**Fig. 5.**
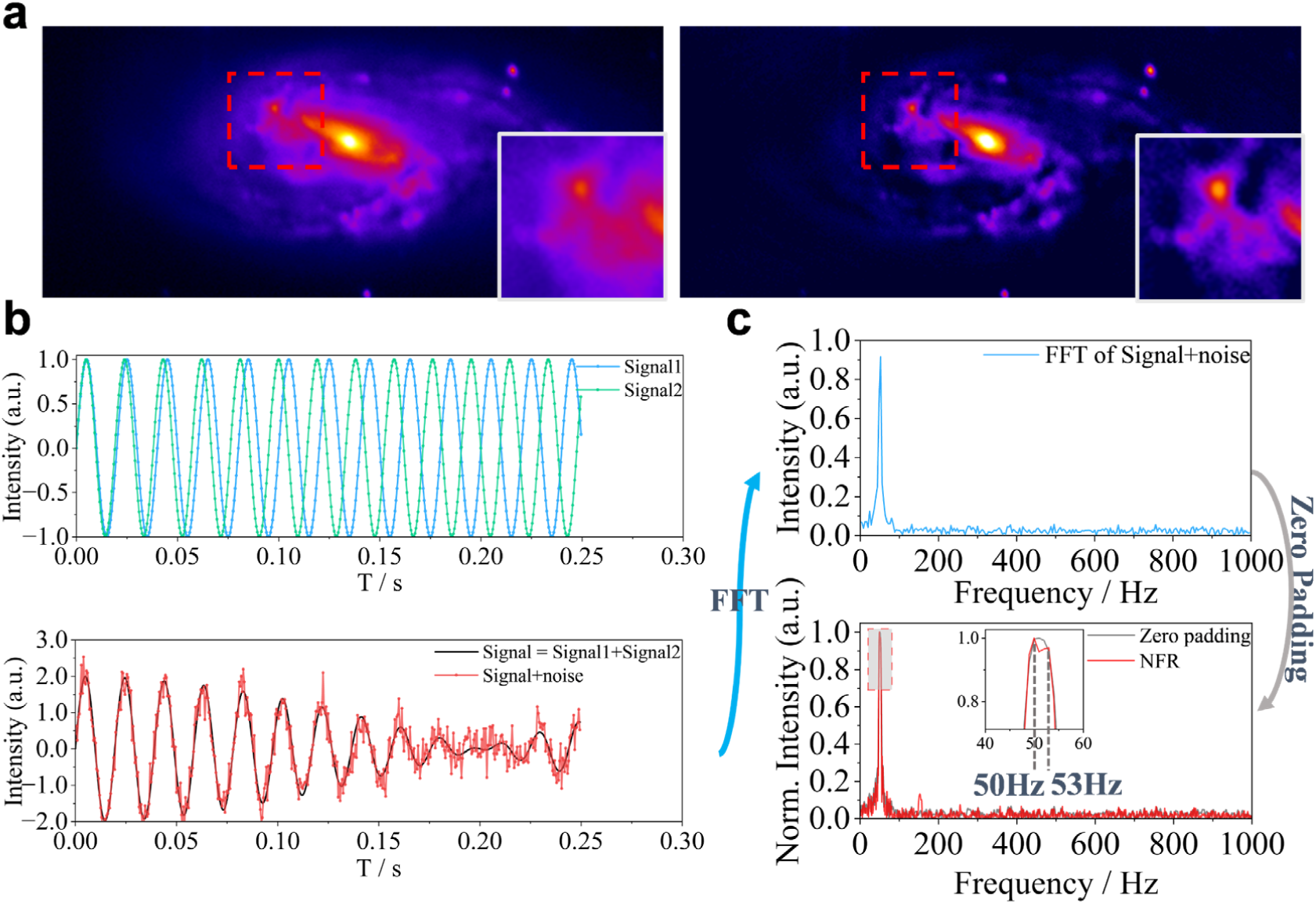
NFR enhances resolution in astronomical imaging and 1D spectral analysis. **a,** NFR enhancement of an image of the M66 galaxy. The processed image (right) shows enhanced detail and contrast compared to the original (left). **b, c** Spectral resolution enhancement of 1D signal. **b**, Generation of a composite signal by summing two closely spaced sinusoids and adding noise. **c,** Spectral analysis showing that a standard Fast Fourier Transform (FFT) (top) and zero-padded FFT (bottom gray line) fail to resolve the two frequency components. NFR (bottom red line) successfully separates the peaks.

More fundamentally, NFR can address resolution limits in one-dimensional signal analysis. A finite observation time T fundamentally limits the achievable spectral resolution to Δf ≈1/T, a consequence of the signal’s true spectrum being convolved with a sinc function^37^. To test NFR’s ability to overcome this limit, we synthesized a 1D signal composed of two closely spaced sinusoids (50 Hz and 52.5 Hz), sampled over 0.25 s (500 points) and corrupted with noise (SNR=20) (Fig. 5b). A direct FFT of this signal reveals only a single, broad peak, as the 2.5 Hz frequency separation is below the theoretical 4 Hz resolution limit (Fig. 5c).

Standard techniques like zero-padding the time-domain signal to 2000 points—which is equivalent to interpolating the spectrum—fail to resolve the two frequencies; they merely smoothen the single composite peak. Remarkably, when NFR is subsequently applied to this interpolated spectrum, it successfully separates the single peak into two distinct components, correctly identifying the constituent frequencies (Fig. 5c). This demonstrates that the NFR principle can achieve a form of spectral super-resolution, resolving frequency components that are inseparable by conventional Fourier analysis due to observation time constraints.

### NFR enhances resolution despite severe optical aberrations

A ubiquitous challenge in practical microscopy is the presence of optical aberrations^38^, which spatially distort the PSF and degrade image quality, particularly when the optical axis is no longer well-aligned (Fig. 6a). While manifesting in diverse forms such as coma and astigmatism, their universal effect is to severely degrade the OTF, disproportionately attenuating high-frequency information. This poses a critical problem for conventional deconvolution, which relies on an accurate, often idealized, and spatially invariant PSF model. When faced with strong off-axis aberrations, the mismatch between the assumed and the real PSF renders deconvolution ineffective.

**Fig. 6.**
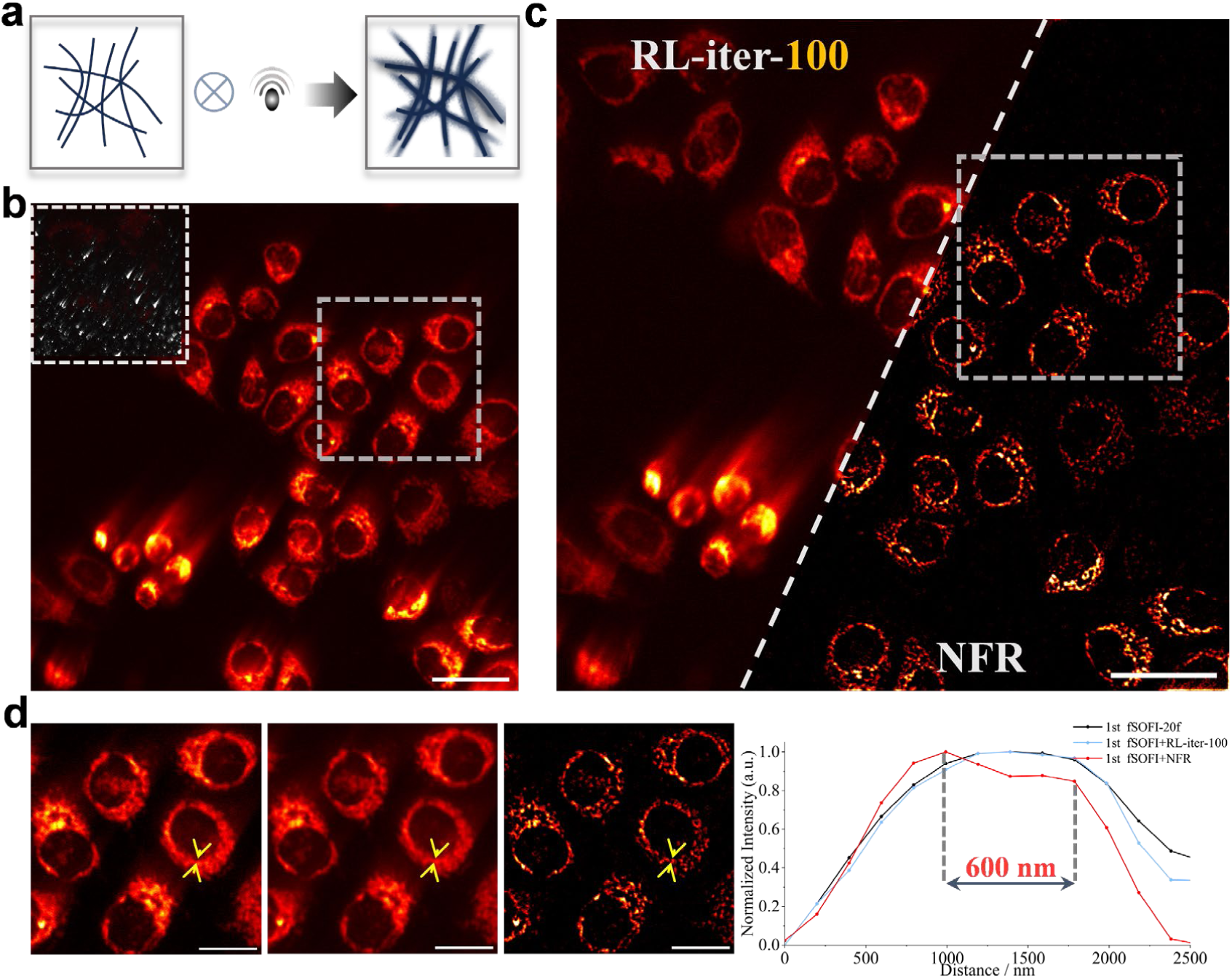
NFR robustly enhances resolution in the presence of strong optical aberrations. **a**, Schematic of image formation with a strong off-axis aberrated PSF. **b**, A two-photon microscopic image of HeLa cells, captured with a system exhibiting strong comatic aberration, as shown by the measured PSF (inset). Scale bar, 40 μm. **c,** Comparison of the aberrated image from **b** processed with Richardson-Lucy (RL) deconvolution and NFR. Scale bar, 40 μm. **d**, Magnified views of the boxed regions from **b, c** and corresponding intensity profiles along the yellow arrow. The profiles demonstrate NFR’s ability to resolve adjacent structures that remain blurred in both the raw and RL-processed images. Scale bar, 20 μm.

To test NFR’s performance under such challenging, real-world conditions, we imaged mitochondria in HeLa cells using a two-photon microscope with significant, uncorrected off-axis aberrations. The raw image exhibits characteristic comet-like streaking and loss of detail, and the distorted, asymmetric PSF clearly indicates the presence of severe coma and astigmatism (Fig. 6b). We then processed the image with both NFR and a 100-iteration RL deconvolution algorithm (Fig. 6c). The RL result shows almost no improvement over the raw data, failing to correct the aberration-induced blur. In striking contrast, a single application of NFR substantially enhances the image, mitigating the visual impact of the aberrations and enhancing structural definition.

Quantitative analysis confirms this superiority. An intensity profile drawn across two adjacent mitochondria (indicated by yellow arrows and detailed in Fig. 6d) reveals that NFR successfully resolves the two structures, separated by approximately 600 nm. Conversely, the profiles for both the raw image (1st fSOFI, equivalent to a 20-frame average) and the RL-deconvolved image show only a single, broad feature, demonstrating their failure to separate the structures. This result highlights NFR’s inherent robustness, as its spectral re-balancing approach does not depend on a predefined, aberration-free system model.

## DISCUSSION

In this work, we introduced the Nonlinear Fourier Re-weighting (NFR) algorithm, a fundamentally new approach to computational image restoration. Traditional deconvolution methods, while powerful, operate on the principle of solving an inverse problem, a process that is inherently dependent on an accurate, often inaccessible, prior model of the system’s PSF. This dependency, coupled with the computational cost and artifact-proneness of iterative solutions, has remained a significant barrier to their widespread, real-time application. NFR circumvents this entire paradigm. Our central aim was to achieve the effect of deconvolution— enhanced resolution and contrast— without being constrained by its procedural and theoretical limitations.

The conceptual leap of NFR lies in reframing the restoration task not as a mathematical inversion, but as a holistic spectral re-balancing. We recognized that the resolution-enhancing effect of deconvolution is to elevate the attenuated high-frequency components to a level more commensurate with the dominant low-frequency components. However, as our results indicate, simply boosting or isolating high frequencies yields unnatural, noisy images. A “healthy” image is defined by the intrinsic balance across its entire spectrum. Inspired by logarithmic compression in sensor signal processing, NFR achieves this re-balancing directly in the Fourier domain. By applying a non-linear logarithmic function, NFR restores a more natural spectral profile, thereby enhancing detail in a single, non-iterative step, without resorting to any predefined imaging models.

Our results demonstrate the remarkable robustness and versatility of this approach. NFR consistently delivered superior performance in a host of challenging, real-world scenarios where traditional methods often fail. Even in the presence of severe off-axis aberrations that render model-based deconvolution ineffective, NFR successfully restored image clarity. Furthermore, it robustly enhanced resolution deep within highly scattering biological tissue in both two-photon and three-photon microscopy, extracting meaningful structural information from extremely low-photon-count data where signal was barely discernible. This proves NFR is not merely a filter, but a powerful information recovery tool.

Beyond its robustness, NFR exhibits profound generality. Its successful application to astronomical imaging and its ability to achieve a form of “spectral super-resolution” in 1D signal analysis underscore that its core principle transcends microscopy. This computational simplicity—requiring only a single forward and inverse FFT—translates directly to near-instantaneous processing speeds. This capability is transformative, enabling not only high-throughput analysis of massive static datasets but also, crucially, the real-time visualization of dynamic biological processes, a domain largely inaccessible to iterative deconvolution algorithms.

Looking forward, the unique properties of NFR open several exciting new frontiers. For instance, in structured illumination microscopy (SIM), the Wiener deconvolution step used in reconstruction is a known source of artifacts^39,40^. Because NFR involves no frequency-domain division, it could serve as a superior, artifact-free replacement for the final reconstruction step in SIM. Furthermore, as the near-infrared-II (NIR-II) fluorescence microscopy gains prominence for its deep-tissue imaging advantages^41^, it faces an inherent trade-off with reduced resolution due to longer wavelengths. NFR presents an ideal computational partner to counteract this resolution loss, maximizing the information yield from these powerful imaging modalities.

Ultimately, NFR represents a paradigm shift in making high-fidelity image restoration accessible, fast, and robust. By decoupling resolution enhancement from system modeling, it empowers researchers to instantly boost the performance of virtually any imaging system. However, NFR is not a panacea and has inherent limitations. The logarithmic re-balancing in the frequency domain compresses the dynamic range in the spatial domain, disrupting the linear relationship between image intensity and fluorophore concentration. Furthermore, NFR is optimized for diffraction-limited scenarios and its efficacy diminishes in cases of severe undersampling or for macro-scale imaging of non-sparse objects with non-sparse emission patterns, where diffraction is not the primary limiting factor. Future work could address these challenges. Exploring adaptive, signal-dependent non-linear mappings could potentially mitigate dynamic range compression while preserving resolution gains. Moreover, integrating NFR with sparse reconstruction frameworks or deep learning models^42^ could extend its applicability to undersampled data and other imaging modalities. Nevertheless, as a tool for real-time enhancement, NFR significantly enhances optical throughput and accelerates the pace of discovery, allowing biologists to resolve complex and transient physiological events that were previously beyond reach.

## Methods

### NFR Principle and Parameter Selection

In an idealized scenario (infinite sampling, no noise), the deconvolution of a fluorescence image can be described by a simple inverse filter in the Fourier domain. This operation is defined as:

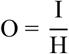

where ***O*** is the restored object spectrum, ***I*** is the measured image spectrum, and ***H*** is the modulation Transfer Function. The objective of this operation is to counteract the low-pass filtering effect of the optical system, which causes a gradual attenuation of signal from low to high spatial frequencies. The ideal result is a restored image spectrum with a uniform energy distribution across the entire passband of the microscope.

The NFR algorithm bypasses the mathematically ill-posed inverse operation of deconvolution. Instead, it aims to directly achieve a spectral energy distribution analogous to that of a successfully deconvolved image. Inspired by the logarithmic response of the human eye to brightness and logarithmic correction in image signal processors, we define the NFR operation as

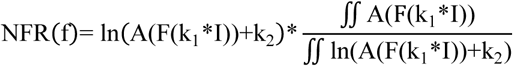

Here, **I** is the input image in the spatial domain, **F** denotes the Fourier transform, **A** is the operation of taking the amplitude of the complex spectrum, and **ln(A(F(k_1_*I))+k_2_)** is the core logarithmic mapping function. The final multiplicative term is a global energy scaling factor that ensures the total energy of the spectrum is conserved before and after the mapping.

The NFR process begins with a FFT of the input image, separating it into its amplitude (A(F(k_1_*I))) and phase spectra ((P(F(k_1_*I))). The NFR operation is applied exclusively to the amplitude spectrum. The core of the operation is the logarithmic function, which nonlinearly remaps the spectral energy. The pre-multiplication factor, derived from the ratio of the total brightness before and after the mapping, serves as a normalization step to restore the overall image intensity.

NFR is governed by two key parameters: a scaling factor ***k_1_*** and a bias ***k_2_***. ***k_1_*** modulates the overall spectral magnitude before FFT. ***k_2_*** acts as an offset that, by leveraging the properties of the logarithm, effectively discards weak frequency components (where NFR(f) < 0). The selection strategy depends on the image’s SNR. For high-SNR images, higher values for both ***k_1_***and ***k_2_*** can be chosen to preserve high-frequency information and achieve aggressive spectral equalization, though care must be taken to avoid potential spectral energy overflow. Conversely, for low-SNR images, lower ***k_1_*** and ***k_2_*** values are preferable to discard high-frequency components overwhelmed by noise and re-balance the remaining, more reliable frequency band.

The selection of ***k_1_***and ***k_2_*** is not an iterative process for convergence, but rather a tuning step to find the optimal balance between resolution enhancement and SNR. A bisection-like parameter search strategy can identify near-optimal parameters within five steps, which can then be directly applied to entire time-series or 3D stacks acquired under identical conditions (Supplementary Figure 1). While other non-linear mappings like Gamma correction can also achieve re-balancing, the logarithmic function is superior due to its inherent capacity for noise suppression (Supplementary Figure 4).

Finally, the NFR process culminates in the reconstruction of the enhanced image. This is achieved by first recombining the NFR-mapped amplitude spectrum (NFR(f)), with any non-positive terms removed, with the original, unmodified phase spectrum (P(F(k_1_*I))). An inverse Fourier transform (iFFT) is then applied to this newly formed complex spectrum. The final NFR-enhanced image is produced after a subsequent automatic contrast adjustment.

### Processing Speed and Implementation

The processing speed of the NFR algorithm was evaluated using its MATLAB implementation. All benchmarks were performed on a standard laptop computer (CPU: Intel Core i7-12700H, RAM: 16 GB) without leveraging GPU acceleration. The time required to process a single 512×512 pixel image was measured to be under 8 ms. For a larger 1024×1024 pixel image, the processing time was under 12 ms.

### NFR Workflow for 1D, Time-Series, and 3D Data

#### One-Dimensional (1D) Signals

For 1D signals, where finite acquisition time limits spectral resolution, NFR is applied directly to the amplitude spectrum of the signal’s Fourier transform to enhance the separation of frequency components.

#### Time-Series Image Sequences

Time-series data are processed frame-by-frame. Optimal NFR parameters are determined from a representative frame and then applied uniformly to the entire sequence, enabling consistent and rapid enhancement.

#### Three-Dimensional (3D) Image Stacks

For 3D image stacks, the workflow is extended by employing a 3D Fast Fourier Transform (***fftn*** and ***ifftn***, MATLAB built-in function). NFR is applied to the resulting 3D amplitude spectrum, typically using parameters determined from a representative central slice. The final volume is reconstructed via a 3D Inverse FFT.

### Microscopy Setups

#### Wide-field Microscopy and Structured Illumination Microscopy (SIM)

Wide-field microscopic and SIM images were acquired using a commercial HIS-SIM system (Guanzhou CSR Biotech Co. Ltd). An Olympus UPLANXAPO 100×/1.50 NA oil-immersion objective was used for all acquisitions. The same excitation and emission settings were used for both microtubule and mitochondrial imaging: samples were excited with a 561 nm laser, and fluorescence was collected through a 609 nm bandpass filter. Images were acquired in both wide-field and 2D-SIM modes. The raw SIM data were subsequently reconstructed using the manufacturer’s Wiener-SIM algorithm.

#### Confocal Microscopy

Confocal microscopic images were acquired on a Zeiss LSM 880 microscope equipped with an Airyscan detector. The pinhole was set to 1 Airy Unit (AU) for all confocal acquisitions. Two objectives were used: a Zeiss Plan-Apochromat 20×/0.8 NA air objective and a Zeiss Plan-Apochromat 40×/1.3 NA oil-immersion objective. For imaging mitochondria, a 561 nm laser was used for excitation, with emission collected through a 609 nm bandpass filter. Airyscan imaging was performed using the system’s integrated 32-channel GaAsP PMT array, which replaces the pinhole. The final image is generated by the ZEN software, which applies a proprietary algorithm that combines pixel reassignment with deconvolution to reconstruct a super-resolution image.

#### Two-Photon Microscopy

Two-photon microscopic imaging was performed on a custom-built microscope based on the Ultima IV platform (Bruker Corporation), coupled to a mode-locked Ti:Sapphire femtosecond laser (Chameleon Ultra II, Coherent Inc.) providing pulses with a <200 fs duration at an 80 MHz repetition rate. Laser power was modulated using Pockels Cells (Conoptics). An Olympus UPLANSAPO 100×/1.40 NA oil-immersion objective was used for all acquisitions. For chloroplast imaging, 950 nm laser pulses were used with power kept below 20 mW (measured after the objective). To record photodamage dynamics, 850 nm pulses were used in conjunction with a resonant scanner, with power below 50 mW. Emitted fluorescence was filtered through a 610/70 nm bandpass filter and detected by a GaAsP PMT (Hamamatsu, H10770).

#### Two-Photon Microscopy with Induced Aberrations

For imaging with strong off-axis aberrations, a commercial Olympus FV1200 microscope was used, coupled to a 1040 nm Yb-doped fiber femtosecond laser. Aberrations were intentionally introduced by tilting a reflection mirror positioned between the galvanometric scanners and the scan lens, thereby misaligning the optical axis. The fluorescence signal was collected through an Olympus XLPLN25XWMP2 (NA 1.05) objective, passed through a 570–625 nm bandpass filter, and subsequently detected by the microscope’s built-in PMT.

### Sample Preparation

#### Microtubule Sample

For microtubule fluorescence imaging, we used commercially prepared COS-7 cell samples (NWCell#4; Nano-Micro Imaging, Tianjin, China). These samples feature COS-7 cells with microtubules specifically labeled for fluorescence microscopy.

#### Live-Cell Labeling of the Mitochondrial Inner Membrane

HeLa cells were cultured in 35-mm glass-bottom dishes (Cellvis, D35-20-1.5-N) for 24 h before being stained with 200 nM PK Mito Orange in serum-free DMEM (30 min, 37°C, 5% CO₂). After two washes, the staining medium was exchanged for prewarmed complete DMEM immediately prior to imaging.

#### Mitochondrial Sample

For mitochondrial fluorescence imaging, we used a commercially available FluoCells™ Prepared Slide #1 (F-36924, Invitrogen™, Thermo Fisher Scientific). This slide contains bovine pulmonary artery endothelial cells with mitochondria specifically labeled with MitoTracker™ Red CMXRos.

#### HeLa Cell Mitochondrial Labeling

HeLa cells were cultured in standard complete medium in a humidified incubator at 37 ℃ with 5% CO_2_. For microscopy, cells were seeded into 35-mm glass-bottomed dishes and allowed to grow to approximately 70-80% confluency. For mitochondrial labeling, a mitochondrial fluorescent probe was added directly to the culture medium of the cells to a final concentration of 2 µM. The cells were then incubated for 2 hours under standard culture conditions. Afterward, the labeling solution was aspirated, and the cells were gently washed twice with pre-warmed phosphate-buffered saline (PBS) to remove background signals. Finally, the cells were immersed in fresh culture medium for two-photon imaging using a water-immersion objective.

#### Arabidopsis Thaliana Leaf Preparation

Leaves from wild-type Arabidopsis thaliana (ecotype Columbia-0) were used for imaging. Seeds were first surface-sterilized with 10% sodium hypochlorite for 5 minutes, followed by three to four rinses with sterile water. The seeds were then vernalized by stratification at 4 ℃ for 3 days before being sown on half-strength Murashige and Skoog (1/2 MS) medium plates. The plates were incubated in a growth chamber at 22 ℃ under a 16-hour light/8-hour dark photoperiod. After 12 days of growth, young leaves were harvested and mounted on a glass slide in a drop of water for observation.

### Performance Metrics

#### FRC Resolution

The effective resolution of the images was quantified using Fourier Ring Correlation. The standard FRC analysis requires two images of the same field of view with independent noise realizations^43^. However, for images where the sampling rate is at least twice the theoretical cutoff frequency (Nyquist-Shannon criterion), a single-image FRC approach can be employed^44^. In this method, a single 2M×2N image is split into four M×N sub-images. These are formed into two pairs, and the FRC is calculated for each pair. The final FRC curve is the average of the two resulting curves. The resolution is determined as the reciprocal of the spatial frequency at which the FRC curve falls below a fixed threshold, typically set to 1/7.

#### Structural Similarity (SSIM) and Contrast-Structure SSIM (CS-SSIM)

The Structural Similarity Index (SSIM) was used to measure the similarity between a processed image x and a ground-truth reference image y. The general SSIM index is a combination of three components: luminance (l), contrast (c), and structure (s). For the default MATLAB implementation, the weighting exponents α, β, and γ are set to 1, resulting in the formula:

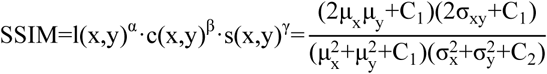

where μ_x_ and μ_y_ are the local means, σ_x_ and σ_y_ are the local standard deviations, and σ_xy_ is the cross-covariance. C_1_, C_2_, and C_3_ are small constants to ensure stability. To specifically evaluate structural and contrast fidelity while ignoring luminance differences, we used the Contrast-Structure SSIM (CS-SSIM). This is achieved by setting α = 0 and β = γ = 1, which simplifies the expression to:

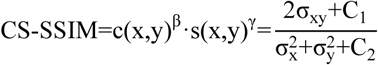

#### Peak Signal-to-Noise Ratio (PSNR)

The Peak Signal-to-Noise Ratio (PSNR) was calculated to quantify the quality of image reconstruction relative to a ground-truth image. It is defined in decibels (dB) as the ratio between the maximum possible power of a signal and the power of corrupting noise. The formula is:

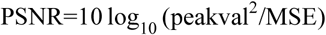

where peakval is the maximum possible pixel value of the image. MSE is the Mean Squared Error between the reference image and the processed image.

### Software Parameters and Simulation

#### Comparative Benchmark Analysis

Images were benchmarked against open-source MATLAB implementations of MRA^29^ and Sparse Deconvolution^14^. For a fair comparison, key parameters (e.g., sparsity, iterations) were carefully optimized for each image to achieve the best possible trade-off between resolution enhancement and artifact suppression.

#### fSOFI Analysis

Fluctuation-based super-resolution imaging (fSOFI) was performed using a MATLAB implementation. The raw image stack was first up-sampled by a factor of two in Fourier space. Subsequently, cumulants of various orders were calculated from the up-sampled temporal data to generate the final fSOFI image.

#### PSF Simulation

All simulations involving a point spread function were based on a theoretical, diffraction-limited PSF. This PSF was modeled as an Airy disk, which was generated using first-kind Bessel functions, assuming incoherent illumination.

#### Richardson-Lucy (RL) Deconvolution

Richardson-Lucy (RL) deconvolution was performed using the built-in ***deconvlucy*** function in MATLAB. The process was controlled by specifying the initial PSF estimate and the total number of iterations as input arguments to the function for blind deconvolution scenarios.

#### NFR Software Implementation

The NFR algorithm was implemented in two primary forms. First, a comprehensive set of MATLAB scripts was developed, providing full functionality for processing 2D images, 3D image stacks, and time-series data. Second, an ImageJ plugin compatible with Fiji was developed, which enables the application of NFR to 2D images and time-series sequences directly within the ImageJ environment.

#### NFR Parameters

The specific NFR processing parameters used for each key image presented in this study are detailed in Table 1. For each image, the table lists the values for the logarithmic mapping parameters, ***k_1_*** and ***k_2_***, as well as the kernel size for the mean filter applied as a post-processing step, where applicable. This information is provided to ensure full reproducibility of our results.

| Figure | $k_1$ | $k_2$ | Mean Filter (radius) |
| --- | --- | --- | --- |
| Figure. 1b c | 1 | 0 | N/A |
| Figure. 2a | 1 | 0 | 0.5 |
| Figure. 2c | 0.75 | 0 | 1.5 |
| Figure. 3 | 0.06 | 0 | 1 |
| Figure. 4a b | 0.2 | 0.6 | N/A |
| Figure. 4d | 0.4 | 0.4 | N/A |
| Figure. 6c | 0.1 | 1 | 1 |
| Extended Data Figure 1.e | 0.006 | 0 | 0 |
| Extended Data Figure 2.b(WF) | 1 | 0.2 | 0 |
| Extended Data Figure 2.b(SIM) | 1 | 1 | 0 |
| Extended Data Figure 3 | 0.1 | 0 | 0 |
| Extended Data Figure 4 | 0.4 | 0 | 1 |
| Extended Data Figure 5 | 0.25 | 0.25 | 1.5 |

## Acknowledgments

We thank Guifeng Xiao from the Core Facilities of Zhejiang University School of Medicine for her valuable assistance with HIS-SIM and LSM 880 imaging; Professor Lin Li from the Institute of Flexible Electronics, Xiamen University, for providing the mitochondrial dye; and Dr. Liang Zhu for his insightful support with the multiphoton microscopic imaging experiments.

---

**Extended Data Figure 1.**
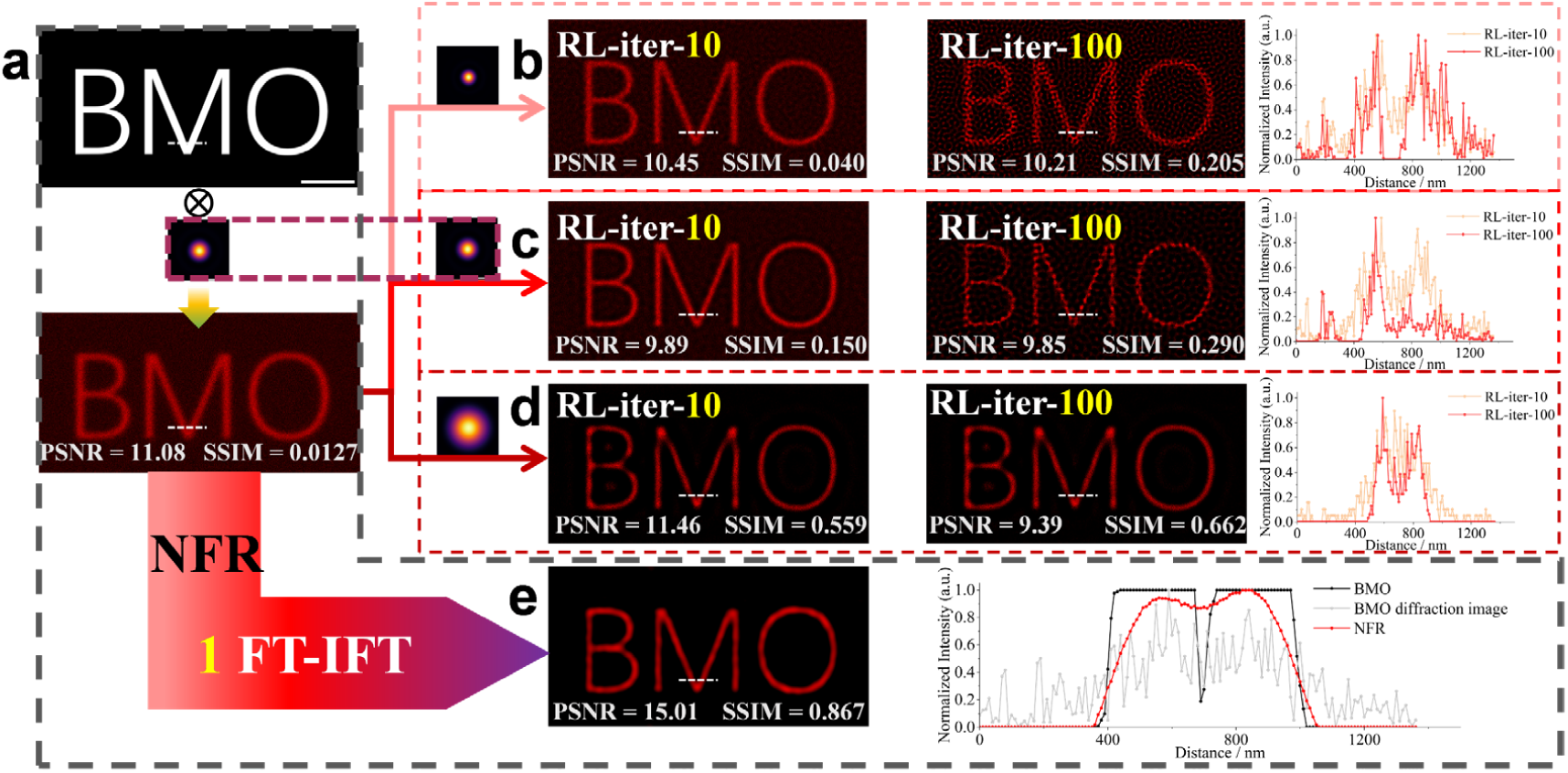
NFR demonstrates superior robustness to noise and PSF mismatch compared to iterative deconvolution. **a,** The noisy, diffracted input image (‘BMO’) (PSNR = 11.08, SSIM = 0.0127) was used for all subsequent processing, compared with the original image without diffraction and noise. Scale Bar, 2 μm. **b-d,** RL deconvolution results after 10 and 100 iterations using **b**, an overly sharp PSF (NA=1.4), **c**, the correct PSF (NA=1.0), and **d**, an overly blurry PSF (NA=0.5). Scale Bar, 2 μm. **e,** The result of a single NFR step applied to the input from **a**. NFR successfully resolves the diffraction-limited ’v’ structure within the letter ’M’ and achieves the highest PSNR and SSIM values among all methods. Scale Bar, 2 μm.

**Extended Data Figure 2.**
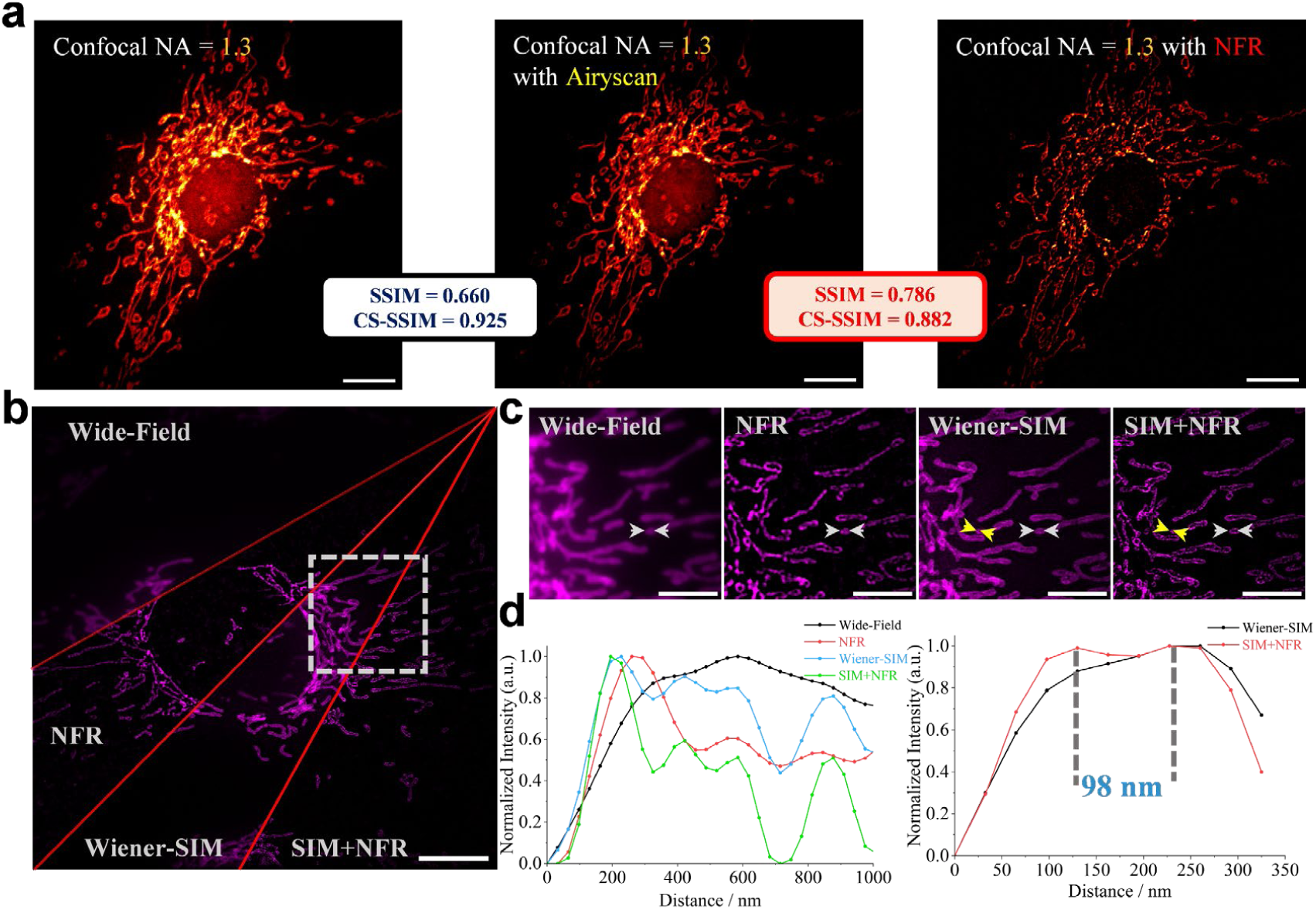
Supplementary validation of NFR performance against Airyscan and on a secondary SIM dataset. **a,** A high-NA (1.3) confocal image and its NFR version are compared to an Airyscan reference. The NFR-enhanced image yields the highest overall SSIM, confirming a high-fidelity enhancement. Scale bar, 10 µm. **b,** A secondary dataset comparing wide-field microscope, wide-field microscope + NFR, Wiener-SIM, and SIM+NFR images of mitochondria. Scale bar, 10 µm. **c,** Magnified views of the boxed region in **b**. Scale bar, 5 µm. **d,** Intensity profiles across mitochondrial cristae (white arrow in **c**) show a progressive resolution gain, with SIM+NFR resolving the finest details unresolved by other methods (yellow arrow in **c**).

**Extended Data Figure 3.**
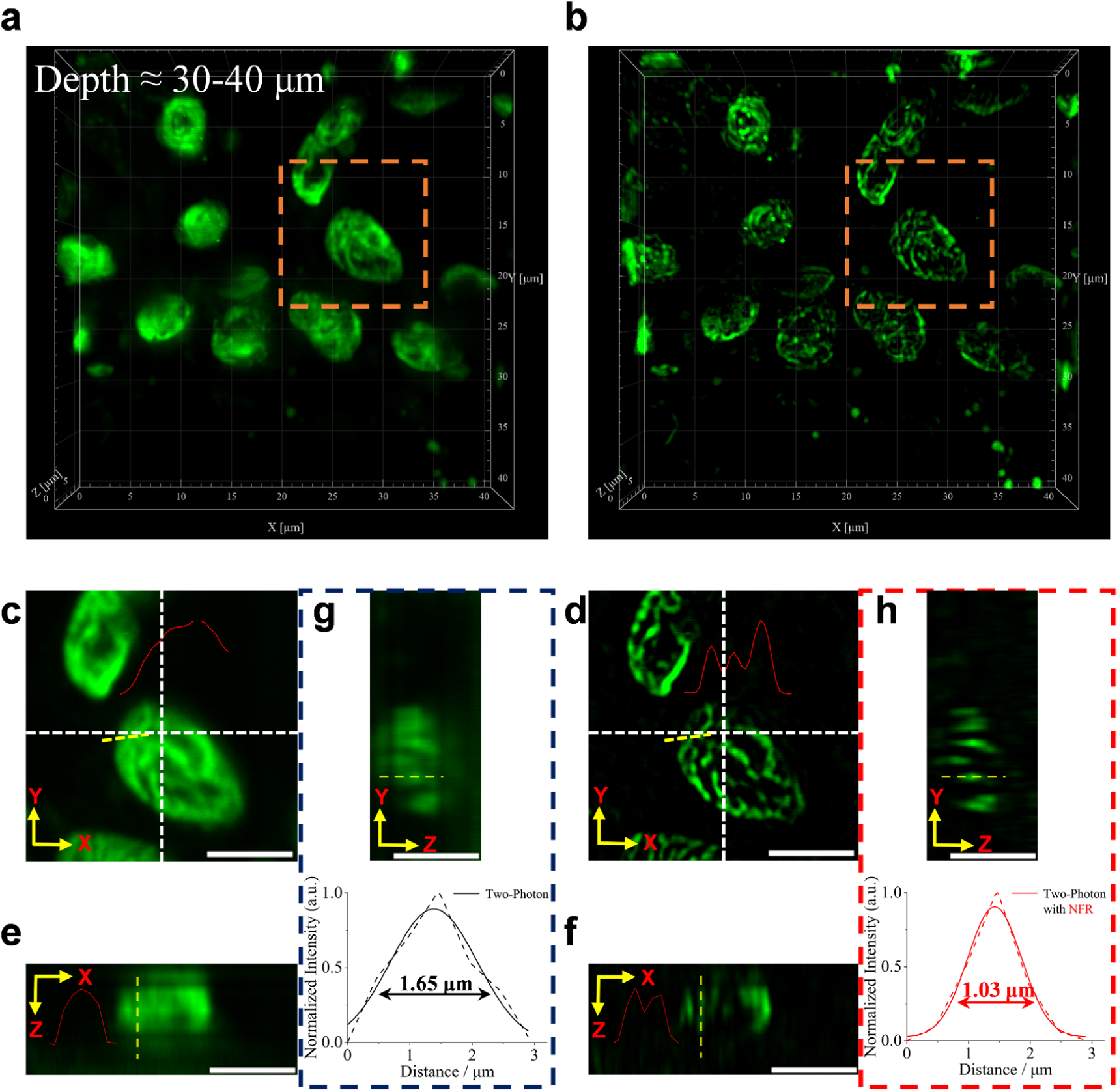
3D-NFR enhances resolution of deep-tissue two-photon microscopic imaging of chloroplasts in a live Arabidopsis leaf. **a,** 3D volume rendering of the raw two-photon microscopic dataset acquired with a voxel size of 79.3 × 79.3 × 500 nm. **b,** The same volume after processing with the 3D-NFR algorithm, showing substantially improved clarity and structural detail. **c, d** Magnified views of the chloroplast indicated by the orange box in **a** and **b**, respectively. The corresponding lateral intensity profiles (XY), red curve, taken along the yellow dashed lines, reveal fine internal structures in the NFR-processed data that are absent in the raw data. Scale Bar, 5 μm. **e, f** Corresponding axial intensity profiles (XZ), red curve, taken along the yellow dashed lines, further confirm that NFR resolves more structural details. Scale Bar, 5 μm. **g, h** Quantitative analysis of a single axial feature (YZ profile) along the yellow dashed line. Gaussian fitting shows that NFR reduces the FWHM from 1.65 µm in the raw data to 1.03 µm, demonstrating a significant enhancement in axial resolution. Scale Bar, 5 μm.

**Extended Data Figure 4.**
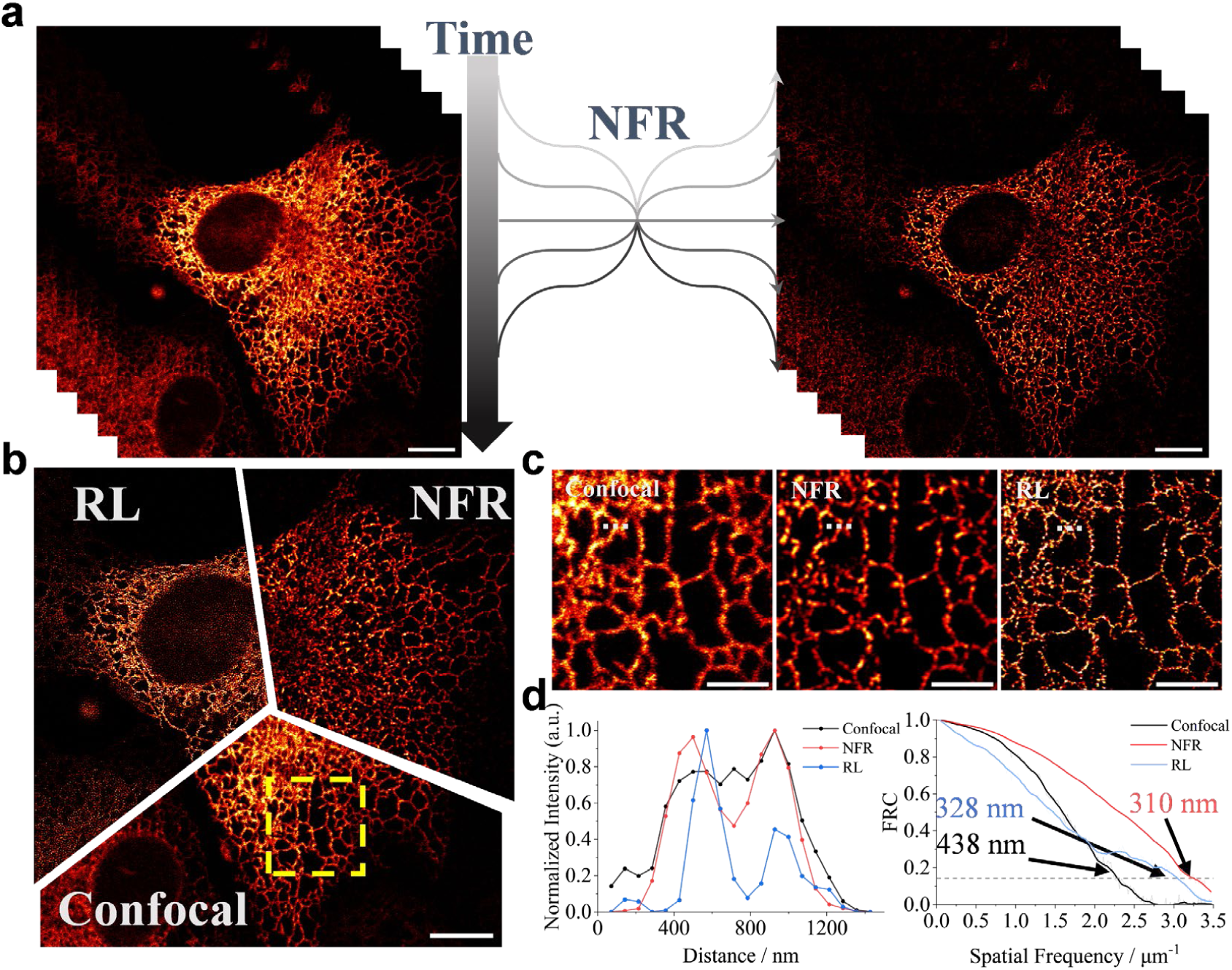
NFR provides high-fidelity enhancement for live-cell time-series^44^ imaging. **a,** Schematic showing the frame-by-frame application of NFR to a time-lapse sequence. Scale bar, 10 μm. **b,** A representative frame comparing the raw confocal image, the NFR-processed result, and the result after 100 iterations of Richardson-Lucy (RL) deconvolution. Scale bar, 10 μm. **c,** Magnified view of the boxed region in **b**, comparing fine structural details across the three methods (left to right: raw, NFR, RL). Scale bar, 4 μm. **d,** Quantitative analysis of the region in **c**. (Left) Intensity profiles along the dashed line in **c**. (Right) Corresponding FRC curves, demonstrating the resolution enhancement achieved by each method.

**Extended Data Figure 5.**
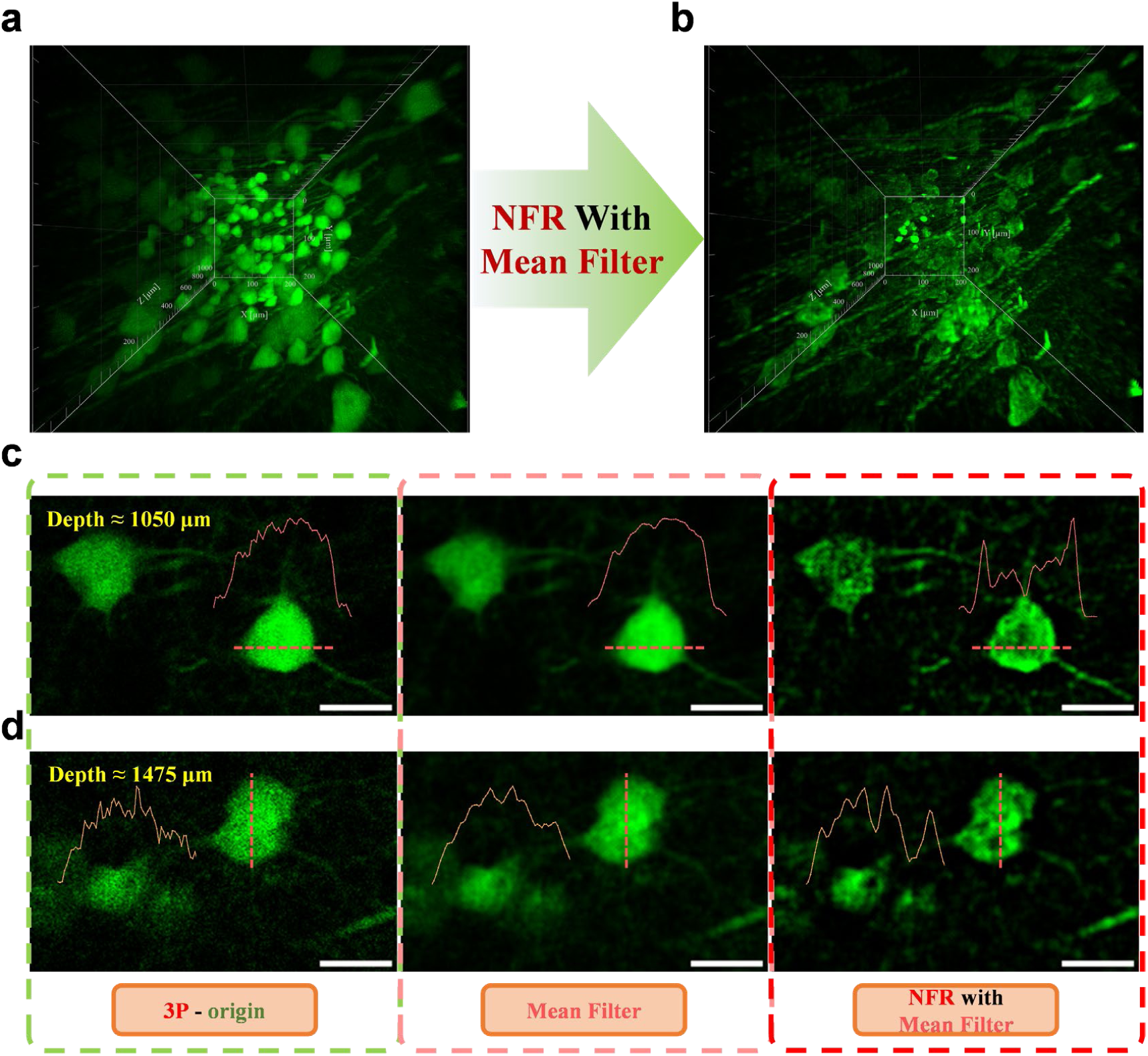
NFR reveals subcellular details in ultra-deep three-photon microscopy. NFR performance on a public three-photon microscopic dataset of YFP-H mouse mPFC. **a, b** Volume renderings of the raw and NFR-processed (post-processed with mean filter) three-photon microscopic stack from a depth of 600-1600 µm. **c, d** XY-slice comparisons at depths of ∼1050 µm and ∼1475 µm. The NFR-processed images (right panels), unlike the raw or mean-filtered data, reveal intracellular structures within neurons, as highlighted by intensity profiles. This demonstrates NFR’s utility in extremely low-photon and highly scattering conditions. Scale Bar, 20 μm.

