## Supplementary Information for "Instant Prior-Free Resolution Enhancement for Cross-Modality Microscopy"

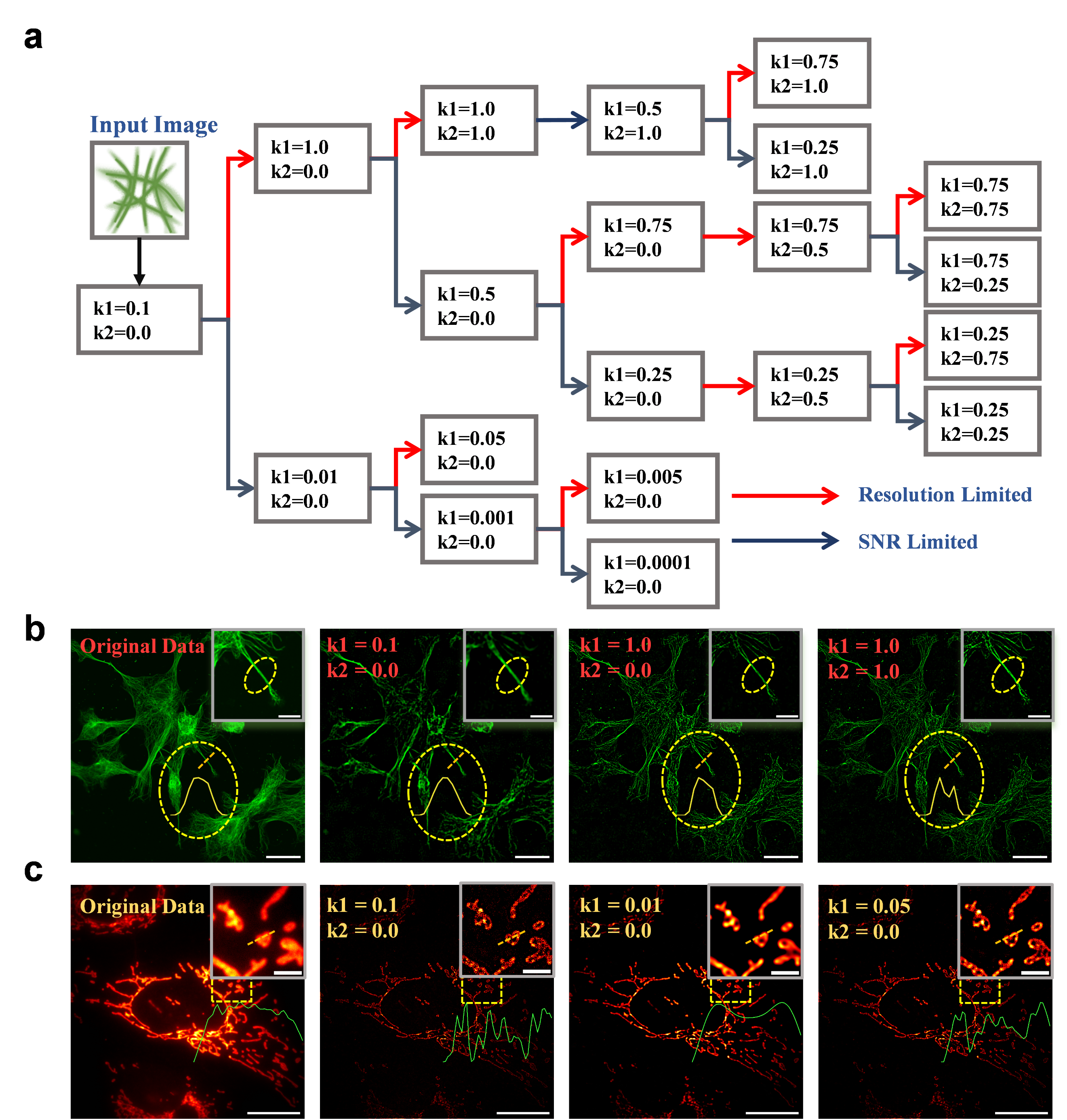
**Supplementary Figure 1 | A systematic strategy for NFR parameter selection to balance resolution and SNR.** **a,** Schematic of the bisection-based workflow for selecting $\text{k}_{\text{1}}$and $\text{k}_{\text{2}}$. **b,** A parameter tuning example on a two-photon microscopic image of microtubule, where three steps are used to achieve maximum resolution and resolve adjacent structures. Scale bars, 30/10 (inset) μm. **c,** A parameter tuning example on a wide-field image of mitochondria, demonstrating a three-step process to find a balanced result between resolution and SNR. Scale bars, 30/5 (inset) μm.

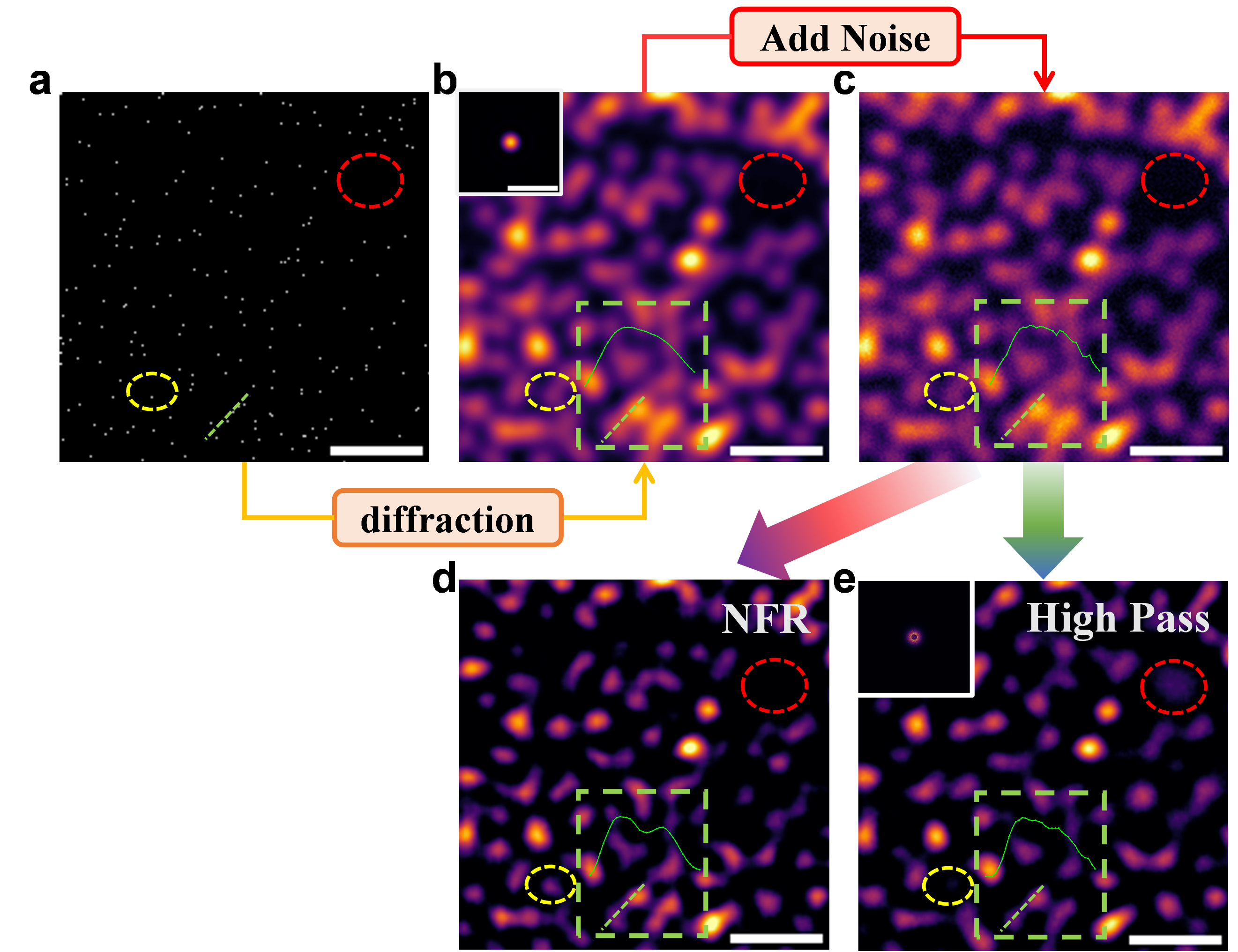
**Supplementary Figure 2| Faithful and robust enhancement of complex structures with NFR. a,** Ground truth object, consisting of randomly distributed point sources. Simulation parameters: NA=1.0, λ=500 nm, pixel size=20 nm. Scale bar: 1 µm. **b,** Ideal diffraction-limited image generated by convolving (a) with the system PSF. Scale bar: 1 µm. **c,** The final input image, created by adding Poisson noise to (b). Scale bar: 1 µm. **d,** The result of processing (c) with NFR, followed by a simple mean filter (radius=1) for noise reduction. Intensity profiles (green box) confirm NFR’s ability to robustly enhance resolution in noisy, multi-point scenarios. Scale bar: 1 µm. **e,** For comparison, the result of processing (c) with a high-pass filter, also followed by a mean filter (radius=1). The high-pass filtered result exhibits both false-positive artifacts (red ellipse) and false-negative information loss (yellow ellipse). In contrast, the NFR result faithfully represents the ground truth. Scale bar: 1 µm.

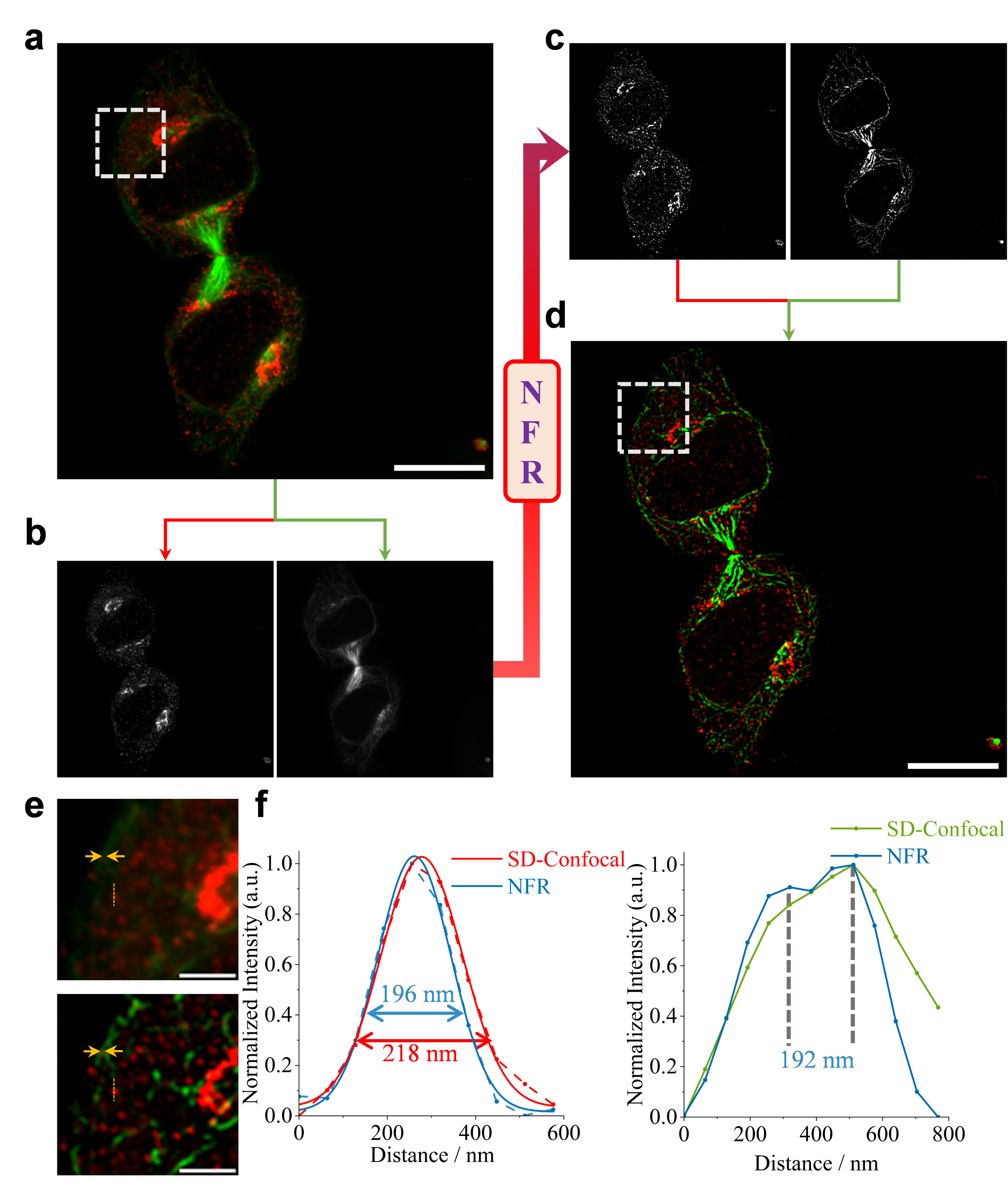
**Supplementary Figure 3| NFR workflow for multi-channel fluorescence imaging. a**, Original two-channel immunofluorescence image of a dividing HeLa cell^1^ (Golgi in red, microtubules in green), Scale bars: 10 µm. **b**, Grayscale images of the separated channels. **c**, Grayscale images after independent NFR processing. **d**, Final NFR-enhanced image after channel merging, Scale bars: 10 µm. **e**, Magnified comparison of regions from the original (top) and NFR-enhanced (bottom) images, Scale bars: 2 µm. **f**, Quantitative analysis demonstrating FWHM reduction in Golgi structures (left, from 218 nm to 196 nm), yellow dashed line in **e**, and improved resolution of adjacent microtubules (right, resolving a 192 nm separation), yellow arrows in **e**.

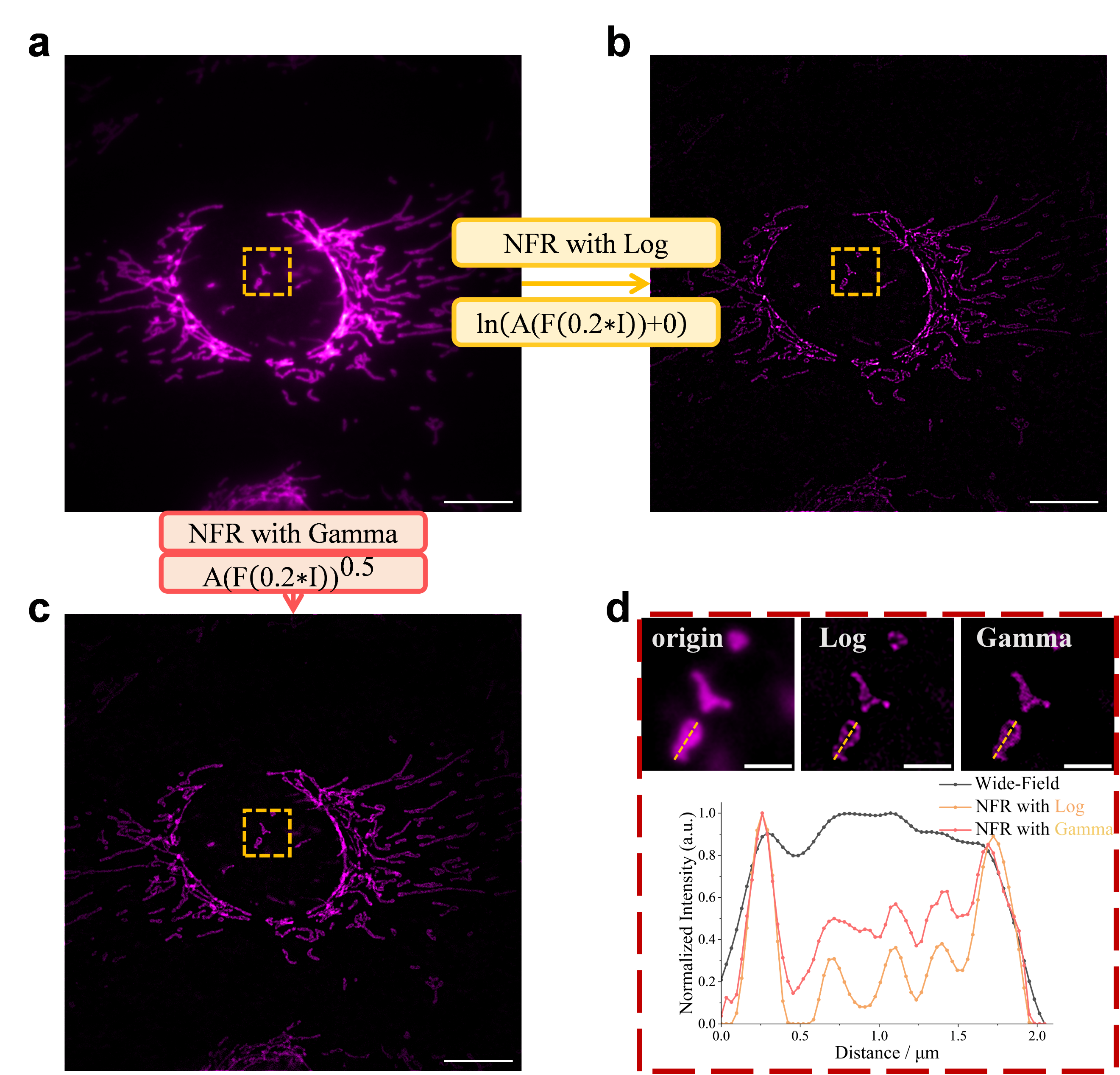
**Supplementary Figure 4| Justification for logarithmic mapping in NFR and validation of the spectral re-balancing principle. a,** A raw wide-field fluorescence microscopic image of mitochondria, where only the outer contours are visible, Scale bar: 10 µm. **b,** The result of processing (a) using NFR's logarithmic mapping, revealing internal cristae structures. Scale bar: 10 µm. **c,** The result of processing (a) using an alternative spectral re-balancing based on gamma mapping. Scale bars: 10 µm. **d,** Magnified views of the yellow boxed region from (a-c) and their corresponding intensity profiles. Scale bars: 2 µm. Intensity profiles demonstrate (yellow dashed line) that while both methods enhance resolution, NFR's logarithmic mapping achieves a superior signal-to-noise ratio, making it the preferred approach for high-fidelity restoration.

**Caption for Supplementary Videos**

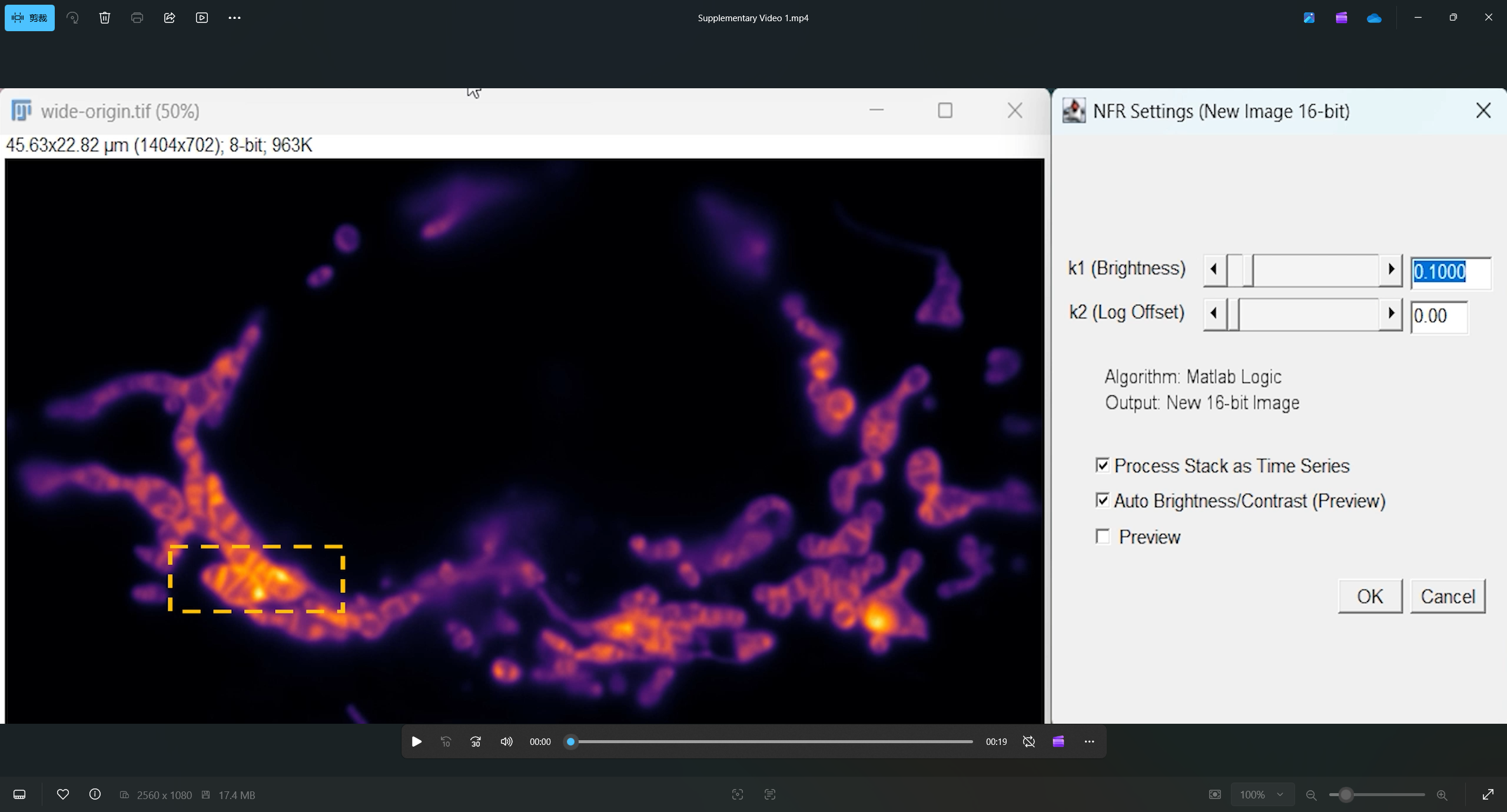
**Supplementary Video 1| Real-time Preview of NFR Enhancement.** This video, captured directly from our custom-developed NFR ImageJ plugin, demonstrates the significant practical advantage of NFR’s non-iterative design for parameter tuning. Leveraging its single-pass nature, which requires only one forward and inverse Fourier transform, NFR enables real-time preview of enhancement results on a standard CPU, without any need for GPU acceleration. The video showcases the enhancement of a mitochondrial inner membrane image as the NFR parameters are adjusted. Specifically, the parameter $\text{k}_{\text{1}}$, which acts as a global scaling factor in the Fourier domain, governs the trade-off between the signal-to-noise ratio (SNR) and the achievable resolution. Smaller values of $\text{k}_{\text{1}}$ result in a smoother image with modest resolution gain, while larger values preserve more fine details at the cost of a lower SNR, which can be easily mitigated with simple denoising steps (e.g., a mean filter) if needed. The primary advantage highlighted here is the near-instantaneous feedback loop, allowing users to intuitively find the optimal parameters for their specific sample. This stands in stark contrast to iterative deconvolution methods, where such real-time exploration is computationally prohibitive.

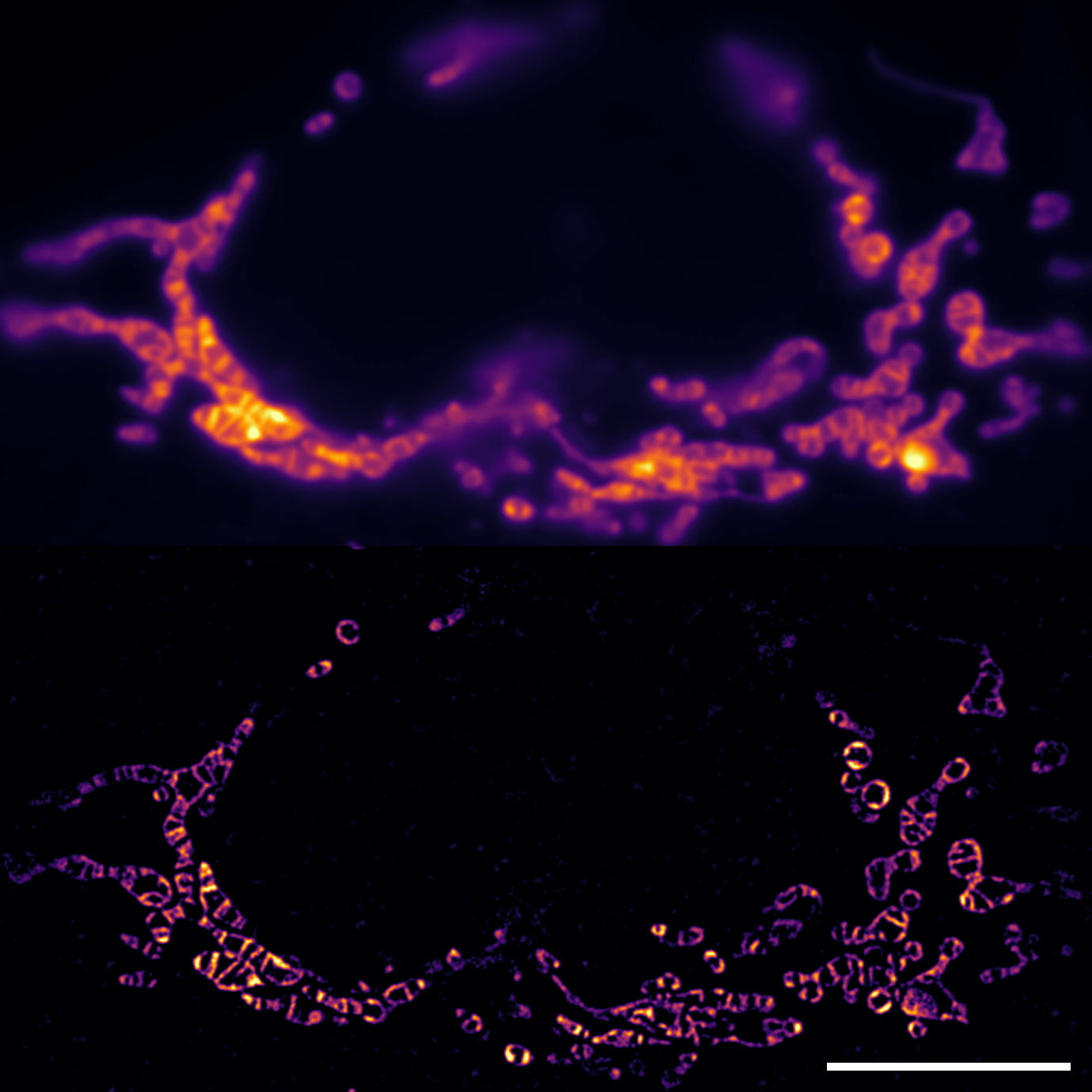

**Supplementary Video 2 | Robust NFR Enhancement of Mitochondrial Inner Membrane Dynamics.** Comparison of a raw widefield time-lapse video (top) of mitochondrial inner membrane dynamics in a live HeLa cell and the corresponding NFR-enhanced result (bottom). The enhancement was performed frame-by-frame using a single, fixed set of NFR parameters, which were interactively determined as demonstrated in our ImageJ plugin (Supplementary Video 1). This highlights NFR's robustness and its suitability for the high-throughput processing of dynamic biological videos, as no frame-specific re-tuning is necessary. Scale bar: 10 µm.

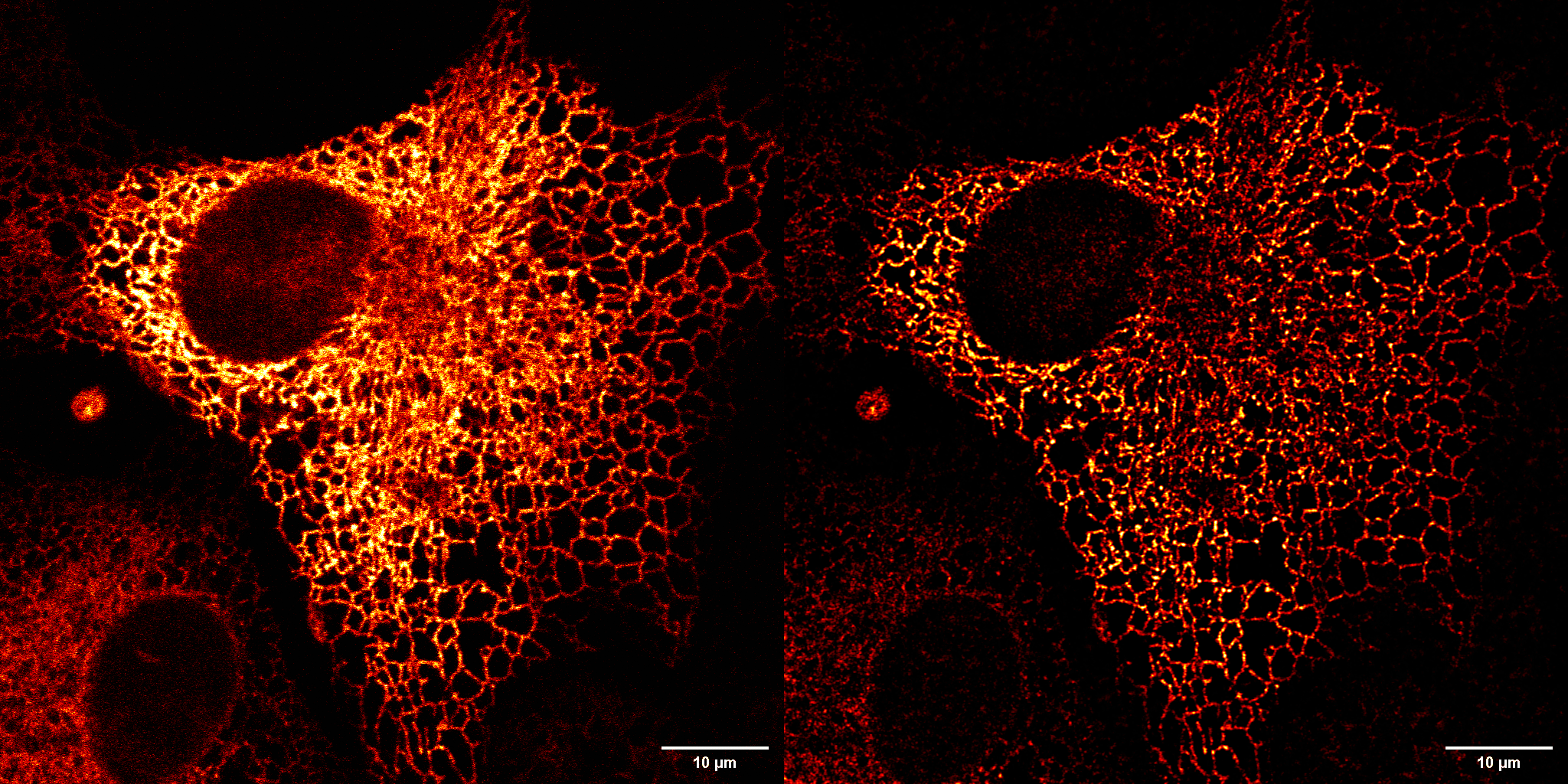

**Supplementary Video 3| NFR enhancement of high-speed endoplasmic reticulum (ER) dynamics.** This video demonstrates the application of NFR to a high-speed time-lapse movie of ER dynamics in a live COS-7 cell expressing Organelle Lights ER-GFP. The data was sourced from a public repository (Cell Image Library, CIL: 723)^2^, a scenario where the system PSF is unknown and cannot be measured, making it ideal for NFR but challenging for deconvolution. The original footage was acquired at 25 frames per second for 100 frames on a Zeiss LSM 510 META. The video displays the raw data (left) alongside the NFR-enhanced result (right). NFR's frame-by-frame processing reveals fine tubular structures and rapid fusion/fission events that are obscured in the raw data, demonstrating its utility for real-time analysis of fast biological processes.

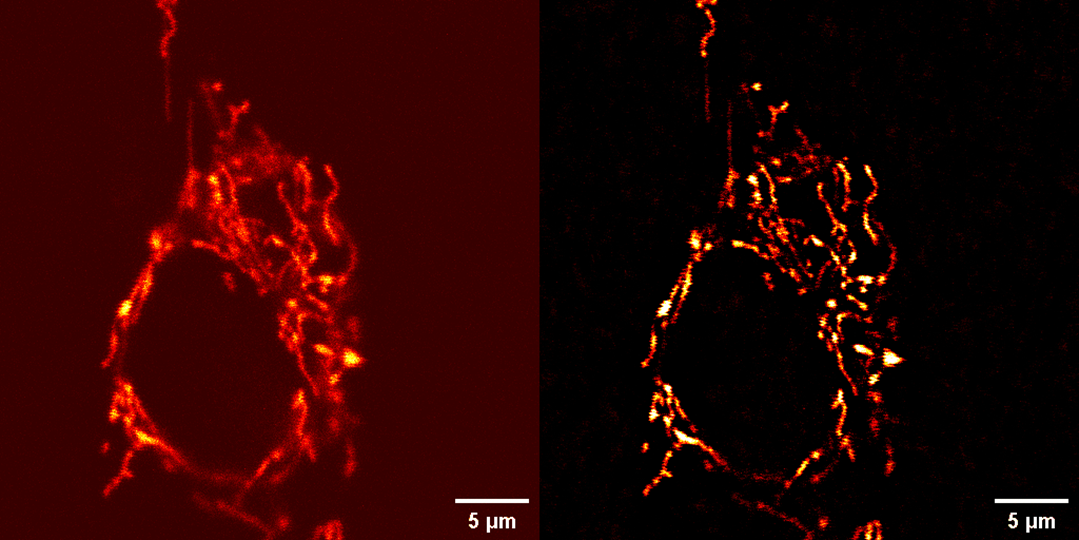

**Supplementary Video 4| NFR enhancement of low-SNR mitochondrial dynamics.** Enhancement of a low signal-to-noise ratio (SNR) time-lapse of mitochondrial dynamics in live NRK cells. The video compares the raw data (left, CIL: 9070, 5.9 s/frame)^3^ with the NFR-processed result (right), demonstrating NFR's ability to improve clarity in noisy datasets without significant noise amplification.

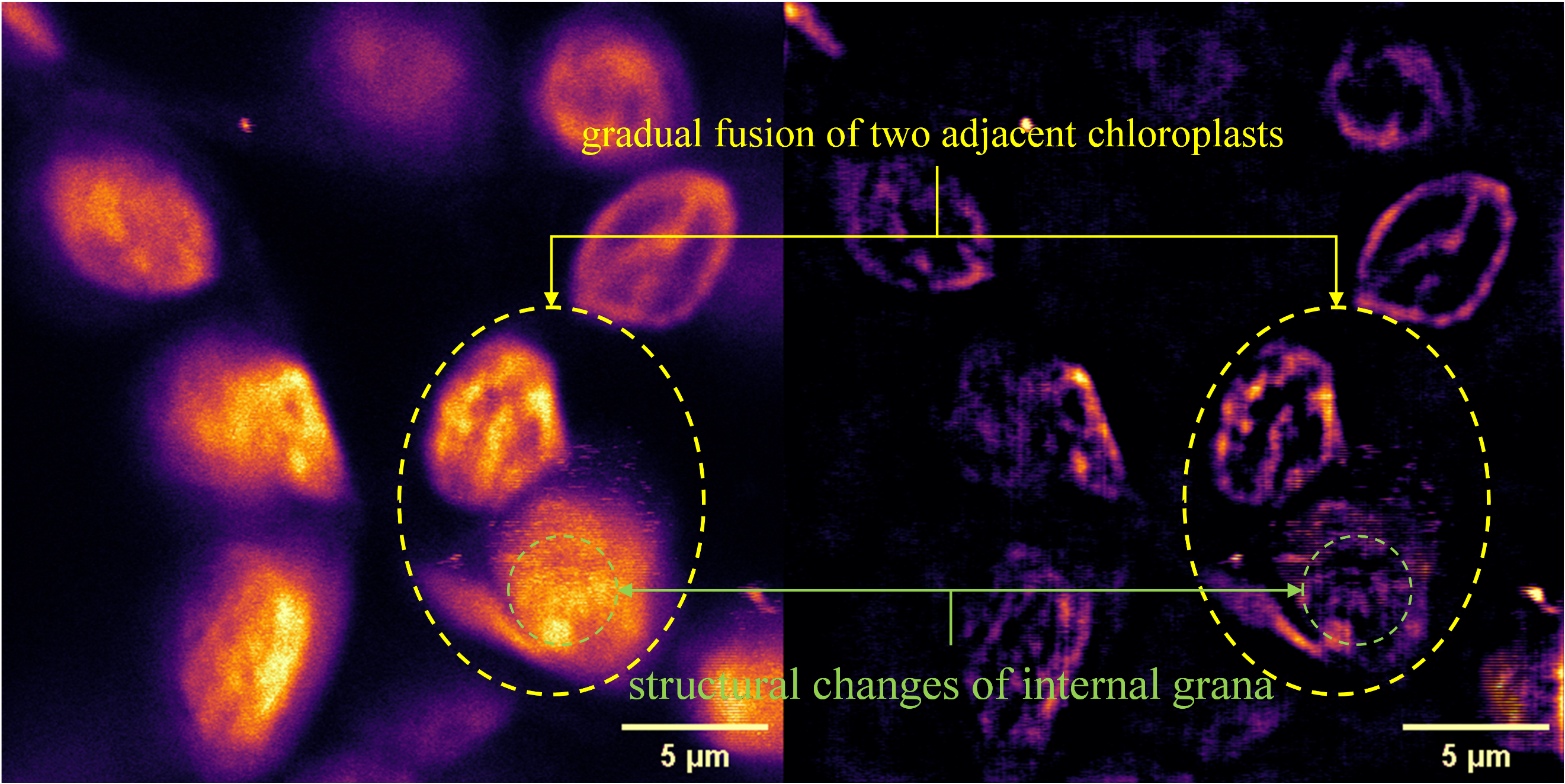
 **Supplementary Video 5| NFR enhancement of chloroplast photodamage.** Real-time enhancement of a two-photon microscopic time-lapse capturing chloroplast photodamage in a live Arabidopsis leaf (850 nm excitation, 3.7 fps). The video compares the raw footage (left) with the NFR-processed result (right), revealing detailed morphological changes and demonstrating NFR's utility for tracking rapid biological events.
